# Substrate-dependent oxidative inactivation of a W-dependent formate dehydrogenase involving selenocysteine displacement

**DOI:** 10.1101/2024.01.10.571421

**Authors:** Guilherme Vilela-Alves, Rita R. Manuel, Aldino Viegas, Philippe Carpentier, Frederic Biaso, Bruno Guigliarelli, Inês A. C. Pereira, Maria João Romão, Cristiano Mota

**Author notes:** (C.M.), (M.J.R.) and (I.A.C.P.).

## Abstract

Metal-dependent formate dehydrogenases are very promising targets for enzyme optimization and design of bio-inspired catalysts for CO_2_ reduction, towards novel strategies for climate change mitigation. For effective application of these enzymes, the catalytic mechanism must be fully understood, and the molecular determinants clarified. Despite numerous studies, several doubts persist, namely regarding the role played by the possible dissociation of the SeCys ligand from the Mo/W active site. Additionally, the O_2_ sensitivity of these enzymes must also be understood as it poses an important obstacle for biotechnological applications. Here we present a combined biochemical, spectroscopic, and structural characterization of *Desulfovibrio vulgaris* FdhAB (*Dv*FdhAB) when exposed to oxygen in the presence of a substrate (formate or CO_2_). This study reveals that O_2_ inactivation is promoted by the presence of either substrate and involves forming a new species in the active site, captured in the crystal structures, where the SeCys ligand is displaced from tungsten coordination and replaced by a dioxygen or peroxide molecule. This new form was reproducibly obtained and supports the conclusion that, although W-*Dv*FdhAB can catalyze the oxidation of formate in the presence of oxygen for some minutes, it gets irreversibly inactivated after prolonged O_2_ exposure in the presence of either substrate. These results reveal that oxidative inactivation does not require reduction of the metal, as widely assumed, as it can also occur in the oxidized state in the presence of CO_2_.

## Introduction

While there is an urgent need for climate change mitigation measures, conversion of atmospheric CO_2_ into added value products entails particularly challenging chemical reactions, due to the inherent stability of this molecule.^1–4^ In this context, the highly active, efficient, and selective metal-dependent Formate Dehydrogenases (Fdh) are very appealing systems that enable harnessing millions of years of natural evolution.^5,6^ As such, there is an increasing interest in developing and deploying these natural enzymes and optimized versions, as well as in creating bio-inspired catalytically active model compounds.^7–12^

Metal-dependent Fdhs harbor at the catalytic site a Mo/W ion coordinating a (seleno)cysteine (SeCys/Cys), four dithiolenes from two Molybdopterin Guanine Dinucleotides (MGD) and one terminal sulfido ligand (-SH/=S). Additionally, [4Fe-4S] clusters are present to convey electrons to/from the physiological partners. While much is already known regarding these enzymes, their catalytic mechanism remains unclear, despite several hypotheses proposed so far. Among these, major divergencies persist, namely whether SeCys has to dissociate from the Mo/W ion during catalysis to allow substrate binding to the W ion^13,14^ or if, instead, a direct hydride transfer occurs between the substrate and the sulfido ligand, with the substrate located in the second coordination sphere.^15^ To date, computational work has addressed several mechanistic hypotheses,^16–19^ but without a clear answer.

Most metal-dependent Fdhs are known to be O_2_-sensitive, easily losing their catalytic activity upon oxygen exposure,^20^ which constitutes a drawback for biotechnological applications. This was addressed for the first time in 1939 by Gale^21^ who described that *Bacterium coli* (now *E. coli*) extracts containing formate dehydrogenase activity react with O_2_ in two different ways, either directly (with O_2_ as electron acceptor in the catalytic cycle) or indirectly, suggesting that this resulted in enzyme inactivation. In 1975, Enoch and Lester^22^ were able to fully reproduce these results, with the isolated enzyme. Additionally, they showed that reduction of dichlorophenolindophenol, ferricyanide, or nitro blue tetrazolium by Fdh was immediately hampered by the presence of O_2_, while complete inactivation of the enzyme required a measure of time, thus restating the role of O_2_ as a competing electron acceptor for Fdh. Remarkably, Graham *et al.*^23^ recently showed that the *Desulfovibrio vulgaris* Hildenborough Fdh2 can oxidize formate in the presence of atmospheric O_2_, unlike most metal-dependent Fdhs, and can also use O_2_ as electron acceptor for formate oxidation, reducing it mostly to H_2_O_2_.

To improve the properties of Fdhs and increase their oxygen tolerance, a detailed structural knowledge of the exact nature of the damage imposed by O_2_ is paramount. This information may spearhead the development of protection strategies and/or ways to prevent or recover activity upon inactivation. Recently, we reported that *D. vulgaris* W/SeCys-FdhAB (*Dv*FdhAB, Fdh1) is irreversibly inactivated by O_2_ in the presence of formate, after a period of time.^24,25^

To better understand the inactivation process, in this work we performed detailed crystallographic studies of *Dv*FdhAB under oxygen exposure along time, in the presence of formate or CO_2_, which were supported by spectroscopic and kinetic studies. These experiments enabled the structural characterization of the oxygen induced damage of *Dv*FdhAB, and the conditions in which it occurs.

## Results and Discussion

### O_2_-inactivation of formate-reduced *Dv*FdhAB involves SeCys192 displacement

Recently, we reported a time-resolved study of the structural changes of *Dv*FdhAB occurring upon formate reduction in anoxic conditions.^26^ A similar approach was employed to gain further insight into the structural changes occurring when the formate-reduced enzyme is exposed to O_2_, thus rendering it inactivated.^24,25^ Since the formate-reduced enzyme is more sensitive to O_2_, formate-treated *Dv*FdhAB crystals were first prepared in the anaerobic chamber (<0.1ppm of O_2_). In a previous work,^25^ we showed that an allosteric disulfide bond (C845-C872) controls the activity of *Dv*FdhAB, with the enzyme being activated by its reduction. Thus, in the current work, the non-activated enzyme (C845-C872 disulfide bond oxidized) was used to slow down the catalytic reaction,^26^ aiming at trapping putative intermediates. The formate-reduced *Dv*FdhAB crystals were flash cooled in liquid nitrogen at different time points after exposure to atmospheric O_2_ (Tables 1 and S1). One of the crystals was cooled before O_2_ exposure (Control_Red) to confirm that the protein was fully reduced at the beginning of the experiment, and in agreement with the formate-reduced structure (PBD_ID: 6SDV)^24^ (Figure S1).

**Table 1.**
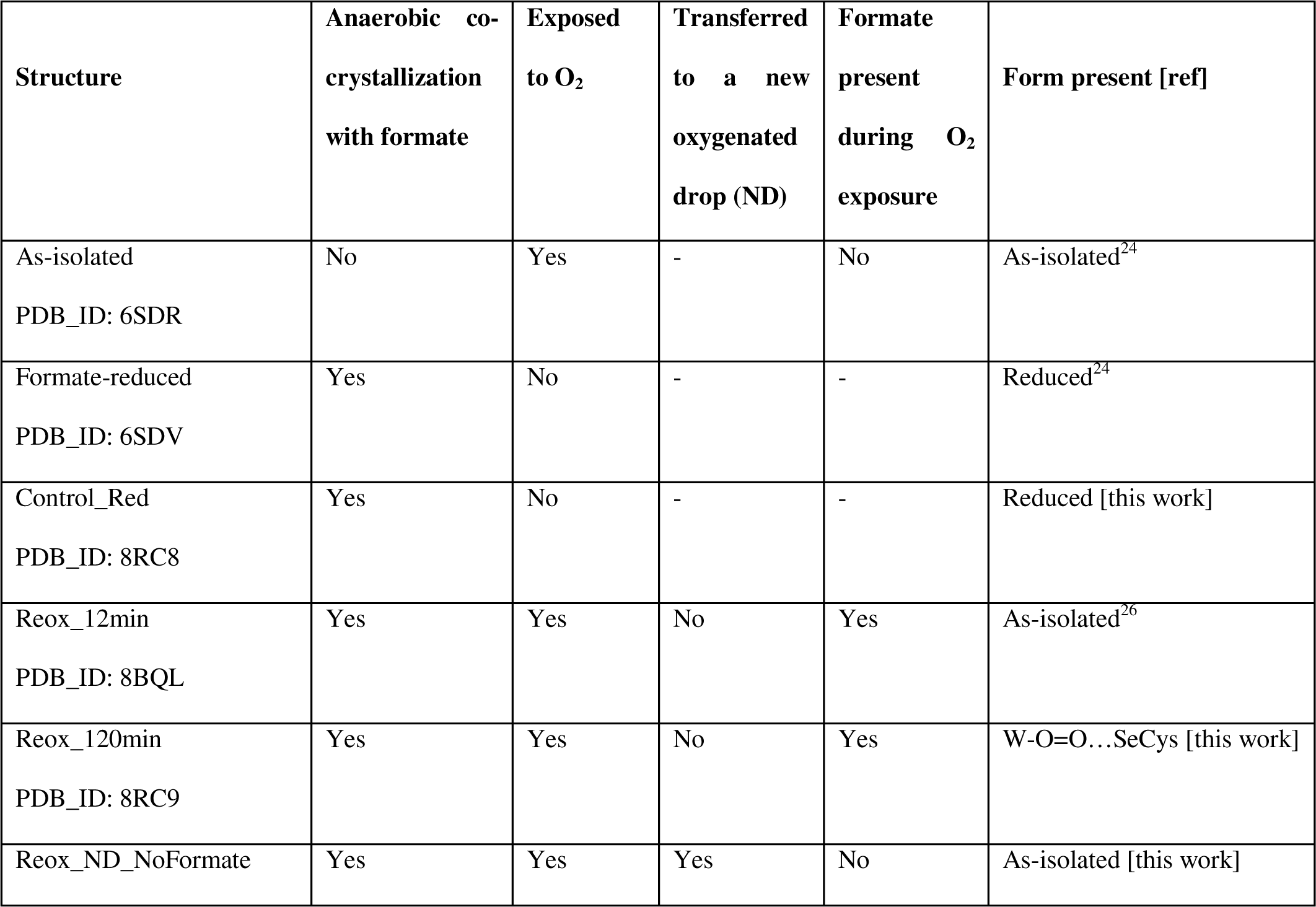
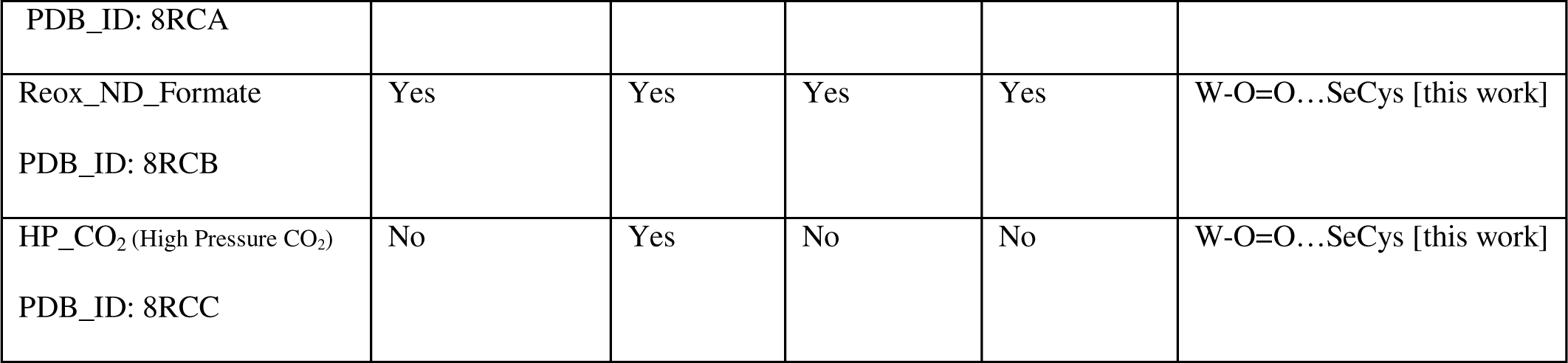
Summary of the different CO_2_ and/or oxygen soaking experiments performed with *Dv*FdhAB in this and other works.

As previously reported,^26^ (PDB_ID: 8BQL), after 12min of O_2_ exposure (Reox_12min) the resulting structure was superimposable with the *Dv*FdhAB as-isolated oxidized form (PDB_ID: 6SDR),^24^ without any sign of damage to the cofactors (Figure S2). However, after 20 minutes of O_2_ exposure, positive and negative electron density close to the active site could be observed. While difficult to interpret, the electron density suggested a mixture of states. After 2 h of O_2_ exposure, the modelled structure (Reox_120min) presented extensive changes in the W active site, when compared to all published structures of *Dv*FdhAB. The most striking difference is the fact that SeCys is no longer coordinated to the W ion, with Se---W distance of 4.2 Å (Figure 1a and 1b, Table S2). The electron density map reveals that the coordination position, that has been left vacant by this displacement, is systematically replaced by a new ligand of diatomic shape in L fashion (W-O bond distances of ∼2.3Å and ∼2.5Å (for the highest resolution structure Reox_120 min (Table S2)). Since O_2_ is expected to react with the active site, this new ligand was interpreted as to be either a dioxygen or a peroxide molecule. Both oxygen atoms are equidistant and within hydrogen bonding distance to the Se atom, suggesting that, once disconnected from the W, the SeCys is in the protonated form. This state of the enzyme was systematically reproduced in multiple experiments (>4 structures). Unambiguous confirmation of the W-Se dissociation is provided by strong anomalous density peaks, detectable at an excess of 11σ, and centered at the positions of the Se atom of SeCys192 and of the W atom (Figure 1a). No trace of anomalous signal was observed at the dioxygen position.

**Figure 1.**
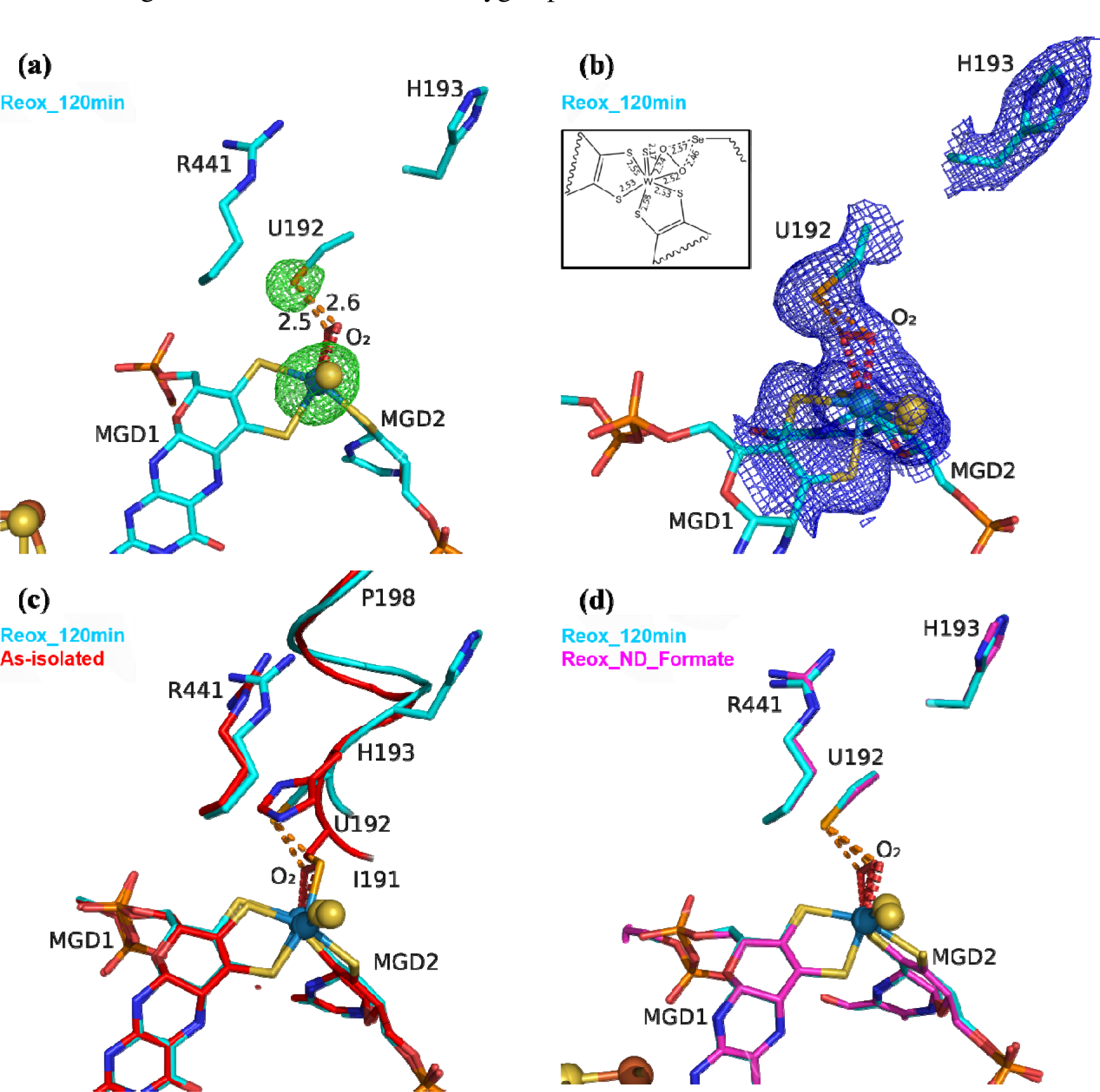
Exposure of formate-reduced *Dv*FdhAB to oxygen leads to SeCys192 displacement. **(a)** *Dv*FdhAB Reox_120min structure (cyan) and corresponding anomalous electron density map contoured at 5σ (green mesh). U192, H193, R441, the two MGD co-factors coordinating the W ion and the dioxygen molecule (red) are shown as sticks. **(b)** Same structure as (a), with the 2Fo-Fc electron density map contoured at 1σ (blue mesh). In the insert, a 2D representation of the W active site indicates the relevant bond lengths (in Å). **(c)** Superposition of *Dv*FdhAB as-isolated (red) and Reox_120min structure (cyan). U192, H193, R441, the two MGD co-factors and the dioxygen molecule (red) are shown as sticks, the helix (I191-P198) is shown as ribbon. **(d)** Superposition of *Dv*FdhAB Reox_120min structure (cyan) and Reox_ND_Formate structure (magenta). U192, H193, R441, the two MGD co-factors and the dioxygen molecule (red) are shown as sticks.

A similar coordination mode of a dioxygen/peroxide molecule to a Mo center has been reported for the *D. gigas (Dg)* AOR (a member of the Xanthine Oxidase family) where this molecule replaces the labile equatorial hydroxyl ligand, being bound to the Mo ion in a ^2^ fashion with Mo-O bond lengths of 1.96 and 2.49 Å for crystals of *Dg*AOR soaked with peroxide (PDB_ID: 4C80, 1.5 Å resolution) (Figure S3).^27^

In the Reox_120min structure, the sulfido ligand and the Se atom were refined with full occupancy revealing B-factors higher than the W and the dithiolene ligands (Table S3). Firstly, this observation suggests that there is an increased uncertainty on the sulfido and Se atom positions due to the reorganization of the W coordination sphere. However, it may also correspond to partial sulfido loss/replacement by an oxygen atom. This same observation is confirmed in the other two structures reported here, with displaced SeCys (W-O=O…Se (see below)) (Table S3).

Along with the dissociation of SeCys192, the helix I191-P198 also changes its position with a screw-like displacement (e.g., 2.9 Å between H193 Cα in as-isolated and Reox_120min) (Figure 1c). Particularly interesting is the large change in H193 that is now facing away from the active site and towards the formate tunnel, with the imidazole ring lodged in the binding position of a glycerol molecule (GOL2) previously reported in several *Dv*FdhAB structures (e.g., PDB_ID: 6SDR) (Figure 1 and Figure S4a). The twisting effect propagates through the α-helix, being attenuated as it moves away from SeCys192. At residues T196/V197 the conformational change is much smaller and comparable with the one between the as-isolated (PDB_ID: 6SDR) and reduced forms (PDB_ID: 6SDV),^24^ and at P198 the structures are essentially superimposable. The displacement of I191 also leads to conformational changes in the sidechain of the hydrophobic core: W533, F160 and F537 (not represented).^24,26^

To overcome the effect of the slow diffusion rate of O_2_ into the anaerobic drop, anaerobic crystals were transferred to a new drop (ND) with similar composition to the mother liquor containing 10 mM of sodium formate, but already oxygenated, and were further exposed to atmospheric oxygen for 34 min (Tables 1 and S1) (a shorter exposure time led to a mixture of states, while a longer one led to poorly diffracting crystals).

The resulting structure (Reox_ND_Formate) was nearly identical to Reox_120min (Figure 1d), showing that the rate of oxygen diffusion plays a significant role in the rate of the SeCys dissociation process. In addition, formate-treated crystals were transferred to a new oxygenated drop without formate and exposed to atmospheric O_2_ for 1 h (Tables 1 and S1) (Reox_ND_NoFormate). The corresponding structure showed the enzyme without any sign of oxygen-induced changes (Figure S4) and the structure superimposes with the as-isolated form (PDB_ID: 6SDR)^24^ with no apparent differences in the active site or [4Fe-4S] centers. This result reveals an essential role of the substrate formate in the SeCys dissociation.

In face of the pronounced structural changes induced by O_2_ and formate, Thermal Shift Assays (TSA) were performed with *Dv*FdhAB in aerobic conditions. These showed a staggering decrease of ∼40°C of the melting temperate (Tm) when formate is present (43.2 °C vs 83.8 °C with and without formate respectively), reflecting a pronounced decrease in protein stability in the presence of formate and O_2_ (Figure S5), probably due to the large conformational changes in the active site.

### Exposure to CO_2_ and O_2_ also inactivates *Dv*FdhAB and involves SeCys192 displacement

To study the effect of O_2_ in the presence of the CO_2_ substrate, High Pressure (HP) gas soaking experiments were performed at the HP Macromolecular Crystallography Lab (HPMX, ESRF, Grenoble, France).^28^ Unexpectedly, crystals from the aerobically isolated oxidized form (prepared in the absence of formate) pressurized with 48 bar of CO_2_ in aerobic conditions resulted in a structure (HP_CO_2_) nearly identical to the Reox_120min structure (Figure 2) (Table S1). The HP_CO_2_ structure shows the same SeCys dissociated form as well as clear electron density for a dioxygen/peroxide molecule coordinated to the W ion. These data show unequivocally that exposure to oxygen in the presence of either substrate results in the W-O=O…SeCys dissociated form, and that, contrary to previous reports,^20,21,25^ tungsten reduction is not a pre-requisite for enzyme inactivation. Remarkably, a CO_2_ molecule is present near the active site, which has never been observed before. The CO_2_ molecule is found close to the W center (4.3 Å away) and the catalytic SeCys (4.0 Å away), being stabilized by van der Waals contacts with the carbonyl of L440 and with the dioxygen ligand, plus a hydrogen bonding interaction to the NH of G442 (Figure 2d). While the CO_2_ molecule is too far from the metal site to be considered a catalytic intermediate, it likely represents a high affinity position along the CO_2_ diffusion pathway from the solvent to the active site. An identical HP experiment was conducted using anaerobic crystals and revealed no discernible displacement of SeCys or any other alterations in the W site. This observation indicates that the displacement of the SeCys is not attributable solely to the applied CO_2_ pressure.

**Figure 2.**
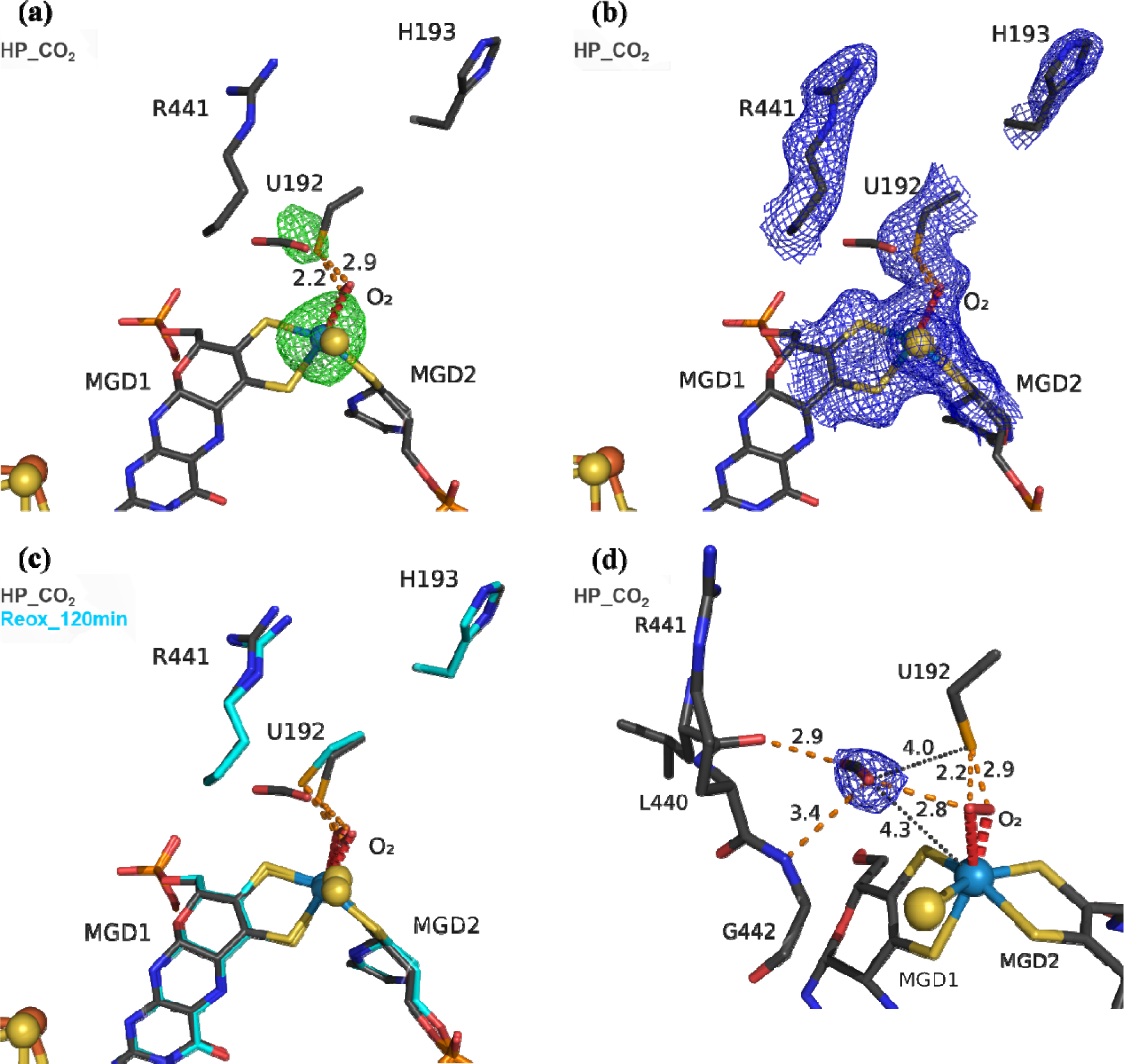
Exposure of *Dv*FdhAB to carbon dioxide and oxygen leads to SeCys192 displacement. **(a)** Structure of *Dv*FdhAB soaked with CO_2_ under High Pressure, aerobically, HP_CO_2_ (dark grey) and respective anomalous electron density map, at 3.5σ (green mesh). U192, H193, R441, a CO_2_ molecule, the two MGD co-factors and the dioxygen molecule (red) coordinating the W ion are shown as sticks. **(b)** Same structure as in (a) with the respective electron density map (2Fo-Fc) contoured at 1σ (blue mesh). **(c)** Superposition of *Dv*FdhAB Reox_120min (cyan) and HP_CO_2_ structures (dark grey). U192, H193, R441, a CO_2_ molecule, the two MGD co-factors and the dioxygen molecule (red) are shown as sticks. **(d)** Same structure as in (a) with the CO_2_ electron density map (2Fo-Fc) contoured at 1σ (blue mesh). The orange dashes represent the contacts between the CO_2_ rear-facing oxygen atom and L440 carbonyl group and the forward-facing oxygen atom and both the G442 NH group and the dioxygen molecule. Additionally, the distances between the CO_2_ forward-facing oxygen atom and the W center and SeCys Se atom are shown as black dotted lines.

The structural data from Reox_ND_NoFormate, Reox_ND_Formate, and the CO_2_ high pressure soaking (HP_CO_2_) clearly demonstrate the simultaneous requirement of oxygen and a substrate for the dissociation of SeCys from the W coordination. When *Dv*FdhAB is oxidized by O_2_ in the absence of formate (Reox_ND_NoFormate), it simply reverts to the as-isolated form and remains in that state, while when formate or CO_2_ are present along with oxygen (Reox_ND_Formate and HP_CO_2_ structures), the dissociated SeCys form emerges, suggesting that the presence of substrate induces a putative intermediate state which progresses to this form.

Since the SeCys dissociated form was also obtained with exposure to oxygen in the presence of CO_2_ (HP_CO_2_), the activity of *Dv*FdhAB was also monitored in these conditions. In fact, co-exposure of a solution of non-activated *Dv*FdhAB to oxygen and CO_2_ led to 80% loss of activity in 30 min (Figure 3a), whereas full inactivation of the DTT-activated enzyme was observed in less than 30 minutes. This behavior is very similar to that observed in the presence of formate and O_2_ as described previously,^25^ where DTT activated *Dv*FdhAB was almost completely inactivated after 30 minutes and completely inactivated after 1h. The control assay, in the presence of CO_2_ in anaerobic conditions, led to a 50% drop in the formate oxidation activity, which is attributed to product inhibition since the CO_2_ reduction activity is not affected in the same conditions (Figure S6). Remarkably, these results confirm that O_2_ inactivation is promoted by the presence of either substrate and not strictly dependent on reduction of the metal, as had been previously assumed.^20,21,25^

**Figure 3.**
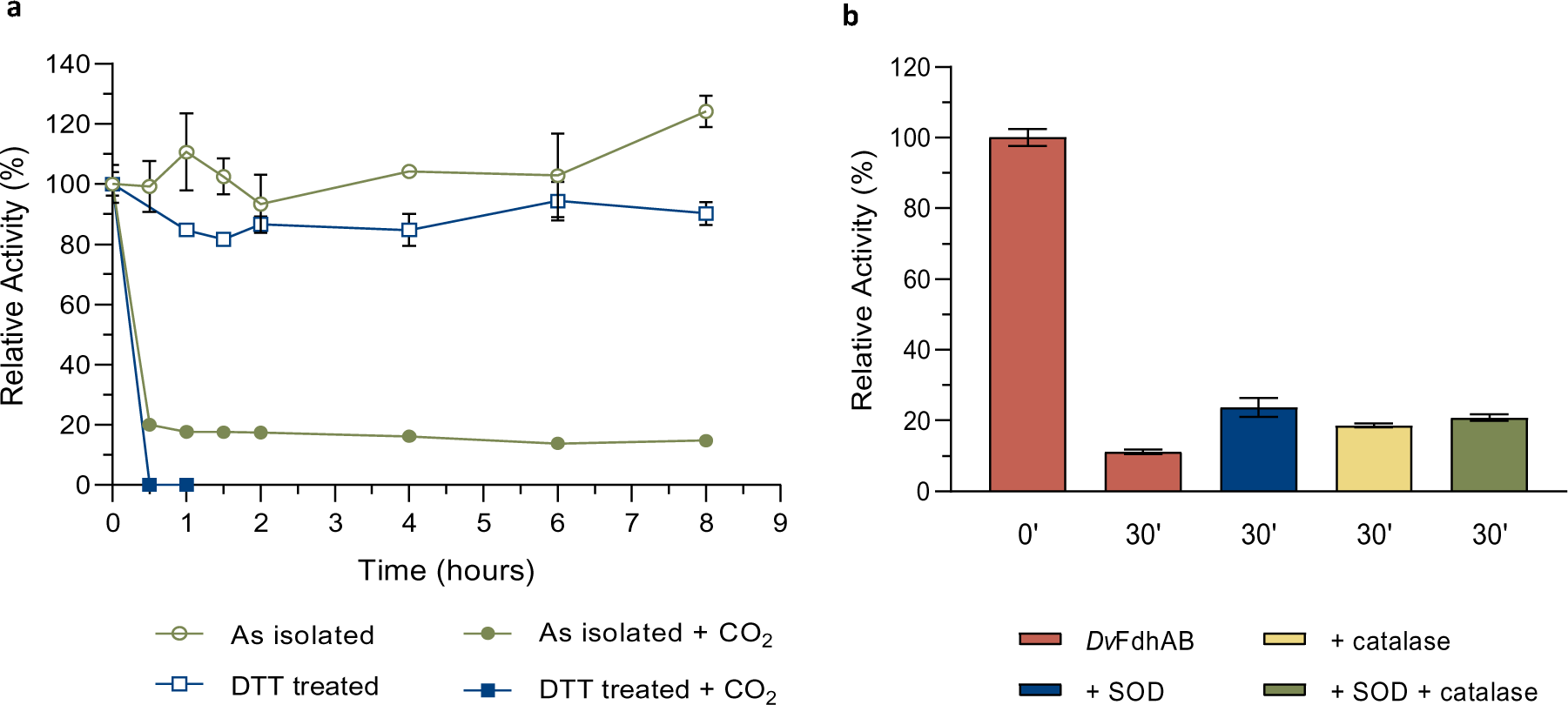
Effect of oxygen on *Dv*FdhAB formate oxidation activity in the presence of substrates. **(a)** Relative activity in the presence of oxygen with the as isolated enzyme (green) and the DTT pre-activated enzyme (blue) in the presence or absence of CO_2_ (100% activity was considered for t=0h; data without CO_2_ are from^24,25^). **(b)** Relative activity in the presence of formate and the ROS scavengers SOD (blue), catalase (yellow) and both (green) versus *Dv*FdhAB alone (red). Data are presented as mean values ± s.d. (n = 3 assay technical replicates).

### *Dv*FdhAB can use O_2_ as electron acceptor

To test if *Dv*FdhAB can oxidize formate using O_2_ as electron acceptor, as reported for the related *Dv* Fdh2,^23^ ^1^H NMR studies were used to analyze formate consumption under aerobic conditions. Using different formate concentrations (1, 10 and 50 mM), formate consumption was indeed observed for some minutes, after which the reaction stops (Figure S7). In the control experiment under anaerobiosis only a very small amount of formate was consumed (≈5% against ≈25% in the presence of O_2_, for 10 mM of formate) (Figure S7). These experiments clearly demonstrate that even in the non-activated form, *Dv*FdhAB can catalyze formate oxidation in the presence of O_2_ for a few minutes, using O_2_ as an electron acceptor. However, for longer O_2_ exposure times, *Dv*FdhAB gets inactivated, in agreement with our reported studies.^24^ The sample recovered from the NMR experiments was concentrated and the protein solution could be crystallized. Although the resulting crystals diffracted at low resolution (2.83 Å), data quality was still good enough to reveal electron density maps at the W active site in agreement with the SeCys displaced form (Figure S8), thus confirming the presence of this species in the inactivated enzyme.

Finally, we measured the formate oxidation activity of *Dv*FdhAB with redox mediators of high redox potential, in the presence and absence of O_2_. The results (Figure S9) show clearly that *Dv*FdhAB has higher activity in the absence of O_2_, in contrast to *Dv*Fdh2,^23^ and confirming its role in anaerobic metabolism.^29^ The activity in aerobic conditions also declines gradually with time, which is reflected in lower reproducibility of the data.

### ROS are not responsible for the substrate dependent oxidative inactivation

The formation of Reactive Oxygen Species (ROS) by metal-dependent Fdhs was reported to be one of the causes of enzyme inactivation leading to the loss of the sulfido group.^20^ In the current work (for all W-O=O…SeCys structures) the sulfido ligand, when refined with full occupancy, has larger B-factors than the surrounding atoms (Table S3), which may also suggest its partial replacement/loss. To investigate if ROS may play a role in the appearance of the SeCys dissociated species, we incubated crystals with different concentrations of hydrogen peroxide (H_2_O_2_). At high hydrogen peroxide concentrations (∼ 10 mM), for different soaking times (1-15 min), the crystals lost diffraction power (no data collection was possible). However, a 10-minute soaking with 1 mM of H_2_O_2_ yielded a 2 Å diffraction dataset and a crystal structure identical to the as-isolated (PDB_ID: 6SDR) structure, indicating that the SeCys dissociation was not promoted under these experimental conditions. Furthermore, ROS protection experiments were performed, by incubating a solution of *Dv*FdhAB with oxygen and formate in the presence and absence of two ROS scavengers (catalase and superoxide dismutase (SOD)). After 30 min of oxygen and formate exposure *Dv*FdhAB activity dropped 90% in the absence of ROS scavengers and 80% in their presence (Figure 3b). This small difference suggests that ROS are not essential to the oxygen inactivation of *Dv*FdhAB but may contribute to it.

### Stabilization of the W(V) state was not possible after SeCys192 displacement

In order to analyze the influence of the structural changes identified by X-ray crystallography on the metal cofactor properties, oxygen and formate exposure assays were also monitored by EPR. Upon anaerobic reduction of the non-activated WT enzyme by formate, the intense EPR signal due to reduced [4Fe-4S]^1+^ centers develops (Figure S10) and the W(V)_F_ signal at g = 1.995, 1.88, 1.85 is detected (Figure 4a). As previously discussed,^25,30^ this W(V) signal represents only 1-2 % of the metal in the non-activated WT enzyme. Upon exposure to aerobic conditions, all these signals vanished (Figure 4b) and at low temperature (Figure S10) an isotropic signal typical of oxidized [3Fe-4S]^1+^ clusters is observed suggesting a partial denaturation of some [4Fe-4S] clusters. Upon degassing to remove O_2_ and further anaerobic reduction with formate, the spectrum of reduced FeS centers is recovered although with a lower intensity. Additional reduction with dithionite led only to a slight increase of this signal, which remained smaller than after the first formate reduction (Figure S10), confirming that a part of the FeS centers was permanently disturbed by oxygen. In contrast, no W(V) signal is recovered after re-reduction with formate or dithionite (Figure 4c and 4d). Similar experiments performed with much longer incubation times with formate and oxygen gave essentially the same results.

**Figure 4.**
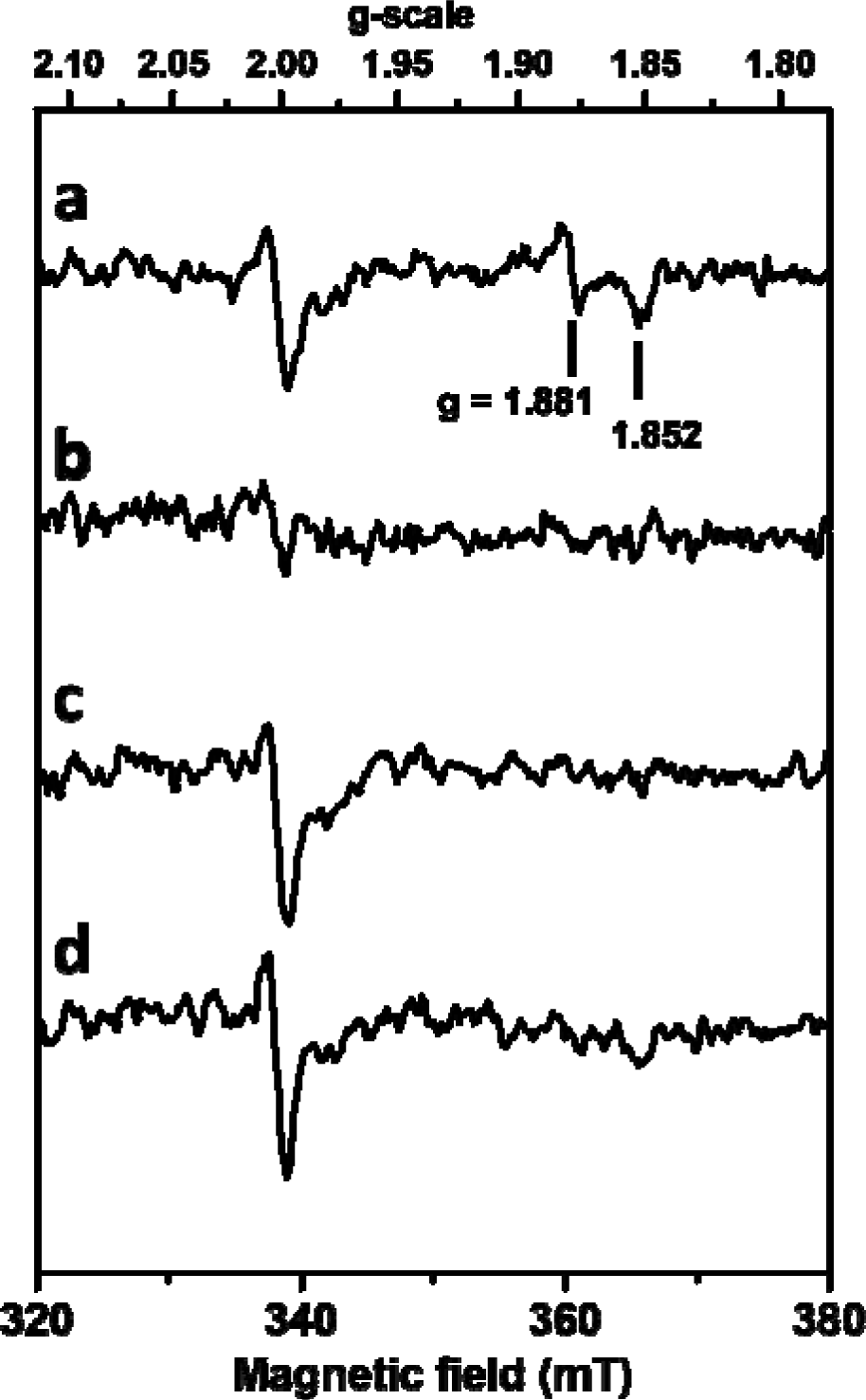
Influence of formate and oxygen exposure on W(V) EPR signals of FdhAB. Anaerobic reduction with formate **(a)** followed by oxygen treatment **(b)**, then degassing and anaerobic reduction with formate **(c)**, and subsequent reduction with dithionite **(d)**. EPR conditions: Temperature, 80 K; microwave power 40 mW at 9.479 GHz, modulation amplitude 1 mT at 100 kHz.

To further evaluate the influence of SeCys displacement on the EPR properties of the tungsten cofactor, a quantum chemistry study was performed on several W(V) cofactor models. Six structural models (numbered 1 to 6) (Figure S11) were built based on the structure of the formate and oxygen-treated *Dv*FdhAB (Reox_120min). These models have in common the coordination of W(V) ion by the four sulfur atoms of the two pterins, a η^2^(O-O) ligand and a displaced SeCys, while they differ in the nature of the exogenous ligand (sulfur or oxygen), the protonation stat of the cofactor and possible H-bonds between the η^2^(O-O) ligand, sulfur and/or selenium atoms (Figure S11). Optimized geometry consistent with protein surrounding constraints could be obtained only for structural models 1-3. The *g-*values of W(V) species were calculated for these models giving values ranging from about *g =* 2.1 to 2.6 (Table S5), much higher than the values observed experimentally for the W(V)_F_ and W(V)_D_ species for the state with bound SeCys ligand.^30^ The lack of any W(V) EPR signals for the enzyme treated with oxygen and formate indicates that the coordination sphere changes experienced by the W atom, notably the SeCys displacement and η^2^(O-O) ligand binding, seem to prevent the thermodynamic stabilization of the W(V) state in the redox potential range investigated.

### Hypothesis for the inactivation mechanism

While the W-O=O…SeCys species reported here was never described before, a SeCys dissociated form of an Fdh was previously observed in the reanalysis of the *E. coli* Fdh-H formate-reduced structure by Raaijmakers & Romao.^13^ However, the low resolution of the data and partial disorder did not enable a proper analysis of the Mo coordination sphere, namely concerning the presence of oxygen or sulfur ligands. In addition, studies on the related *Rhodobacter capsulatus* Mo/Cys-containing Fdh,^31,32^ suggested that the Cys ligand is displaced during catalysis being replaced by an oxygen atom coordinating the Mo. This proposal was based on X-ray absorption spectroscopy studies at the Mo K-edge and NMR experiments with labelled substrates. Interestingly, these data were obtained for a formate-reduced anaerobic sample.

The data now reported raise several hypotheses regarding the SeCys displacement. Formate oxidation in the presence of O_2_ may lead to the production of ROS, which may attack the metal site leading to the SeCys displacement. This is likely to occur in the initial minutes when formate is being oxidized with O_2_ as electron acceptor but cannot explain inactivation in the presence of CO_2_, where no reductant is present. Furthermore, the experiments with ROS scavengers suggest that ROS are not essential to enzyme inactivation, although they may also contribute. A second hypothesis is that the presence of formate or CO_2_ leads to an intermediate conformational state that is more susceptible to oxidative attack, resulting in SeCys dissociation. A third hypothesis is that the dissociated form is a putative catalytic intermediate, which is trapped in the presence of oxygen. However, if the dissociated form was a catalytic intermediate, one would expect the coordination vacancy to be filled with substrate, while the SeCys192 would reoccupy its coordination position upon product release. However, there is not enough evidence to corroborate the third hypothesis, which is also challenged by the multiple time resolved and ligand soakings/co-crystallization studies previously reported^26^ that did not show SeCys dissociation in the absence of O_2_. Therefore, the most likely possibility is that the W-O=O…SeCys dissociated form results from a substrate-induced state where the SeCys-W coordination is more prone to attack by O_2_. The combination of the first and second hypotheses seems more coherent and can explain why this state was only captured in the experiments with oxygen and substrate, while accounting for the role of ROS. The W-O=O…SeCys form seems to correspond to an irreversibly inactivated state since after its formation, the enzyme could not be reactivated by removal of O_2_, DTT treatment or electrochemically by application of a low redox potential.^25^

The oxygen sensitivity of *Dv*FdhAB in the presence of substrate is in line with its protection mechanism involving an allosteric redox switch, which prevents formate binding and catalysis during transient O_2_ exposure, with *in vivo* formate concentrations in the low µM range.^25^ The current study was performed using a formate concentration of 10 mM, which is 5 times higher than the K_M_ of *Dv*FdhAB in the “protected form” (≈2 mM).^25^ Thus, under these conditions formate reduction and O_2_-dependent inactivation is expected to occur.

## Conclusion

In conclusion, the current study shows that the SeCys ligand dissociates from the W center when *Dv*FdhAB is exposed simultaneously to oxygen and one of the substrates (formate or CO_2_), while the vacant position is occupied by a peroxide or dioxygen molecule. This form likely corresponds to a substrate-dependent oxygen/ROS inactivated form, and thus reveals a mechanism for O_2_-inactivation of metal-dependent Fdhs. O_2_ inactivation may also involve loss/replacement of the sulfido ligand as previously reported,^20^ including in *Dv*FdhAB.^25^ These results are also in line with previous reports on the *R capsulatus* Mo-CysFdh with the presence of oxygen ligands bound to Mo, whereby Mo-O(=O) forms accompany Cys ligand dissociation.^31,32^ Regarding the divergences about the possible dissociation of the (Se)Cys ligand during catalysis, we have now demonstrated beyond doubt that this displacement does occur under turnover conditions in the presence of oxygen. The evidence so far suggests that this form most likely corresponds to an inactivation process, although its involvement in the catalytic mechanism cannot be excluded. A deeper understanding of this inactivation process and its possible relation to the catalytic mechanism of Mo/W Fdh enzymes will only be feasible using time-resolved crystallography techniques, supported by spectroscopic data. Such information will be essential in future efforts to engineer an enzyme envisaging the prevention of the substrate/oxygen inactivation, towards continuous catalysis under atmospheric conditions.

## Experimental Section

### Expression and Purification of *D. vulgaris* FdhAB

*Dv*FdhAB was expressed and affinity-purified from *D. vulgaris* Hildenborough, as previously described,^24^ with some modifications. Affinity chromatography was performed with Strep-TactinTMXT 4FlowTM resin (IBA Lifesciences), and the protein eluted with 100 mM Tris/HCl buffer containing 150 mM NaCl and 50 mM biotin. Protein concentration was routinely determined based on E == 43.45 mM^-1^cm^-1^ and purity of samples was judged by 12% SDS-polyacrylamide gel.

### Crystallization, Data Collection, Structure Solution, and Refinement

Crystallization of purified wild type (WT) *Dv*FdhAB was performed in an anaerobic chamber under an argon atmosphere at <0.1 ppm of oxygen, and all the solutions were previously degassed and stored in the anaerobic chamber. All crystals were obtained using the hanging-drop vapor diffusion method, from drops of 2 µL (1:1, protein:precipitant ratio) in 24 well plates (24 well XRL plate Molecular Dimensions) at 20°C. *Dv*FdhAB WT at 10 mg/mL was crystallized in conditions with 22 to 26 % PEG 3350 (w/v), 0.1 M Tris-HCl pH 8.0 and 1 M LiCl, and cocrystallized with 10 mM of sodium formate^24^ with the addition of 0.2 µL of a dilution 1:500 from a stock of microseeds of WT *Dv*FdhAB to the drop (crystals appeared within 24 h and grew during one week).

The crystallization plates were removed from the anaerobic chamber, the wells were opened and atmospheric O_2_ was let to diffuse into the drops for different amounts of time (0 min, 12 min and 2 h) (Tables 1 and S1).

Additionally, two control experiments were performed. Prior to oxygen exposure, crystals were transferred to two aerobic drops of harvesting solution (28 % PEG 3350 (w/v), 0.1 M Tris-HCl pH 8.0 and 1 M LiCl), one with 10 mM of sodium formate and the other without sodium formate. These crystals were then further exposed to atmospheric oxygen for 34 min and 1 h, respectively. After oxygen exposure all crystals were transferred to a cryoprotectant solution consisting of the harvesting solution supplemented with 20% (v/v) glycerol, and then flash cooled in liquid nitrogen.

For the High-Pressure (HP) CO_2_ experiments, crystals were obtained aerobically using the hanging-drop vapor diffusion method, from drops of 2 µL (1:1, protein:precipitant ratio) in 24 well plates (24 well XRL plate Molecular Dimensions) at 20°C. *Dv*FdhAB WT at 10 mg/mL was crystallized in conditions with 22 to 26 % PEG 3350 (w/v), 0.1 M Tris-HCl pH 8.0 and 1 M LiCl^24^ with the addition of 0.2 µL of a dilution 1:500 from a stock of microseeds of *Dv*FdhAB WT to the drop (crystals appeared within 24 h and grew in three days). CO_2_ derivatives were prepared at the ESRF HPMX laboratory.^28,33^ The crystals were pressurized and flash-cooled in a pressure cell in two stages. First, the crystals were soaked in a pressurized atmosphere at 48 bar of CO_2_ pressure from the derivative, at room temperature, then transferred into a helium atmosphere at 48 bar at 77 K to flash-cool and stabilize the derivative. The crystals were thereafter recovered and handled in liquid nitrogen to preserve the CO_2_ derivative state.

X-ray diffraction experiments were performed at the ESRF synchrotron (on beamlines ID23-1, ID30A-3 and ID30B)^34–36^ and the data were processed with either the programs XDS^37^ and Aimless^38^ or autoPROC^39^ and Staraniso.^40^ The Staraniso software was used when data presented anisotropy, to improve overall quality of the final electron density maps. The structures were solved by molecular replacement with Phaser^41^ from the CCP4 suite,^42^ using as search model the previously published formate reduced structure (PDB ID: 6SDV), except for the CO_2_ derivative (HP_CO_2_ structure), for which the as-isolated structure (PDB ID: 6SDR) was used. The models were refined with iterative cycles of manual model building with Coot^43^ and refinement with REFMAC5.^44^ The models were rebuilt with PDBredo^45^ and images produced with PyMOL^46^. Data processing and refinement statistics are presented in Table S4.

### Activity assays

Activity assays for formate oxidation and CO_2_ reduction were measured as previously described.^24^ Assays of oxygen exposure in the presence of CO_2_ were performed with WT *Dv*FdhAB and with DTT-activated *Dv*FdhAB. DTT activation was performed under anaerobic conditions by incubating the enzyme with 50 mM DTT for 2.5 min, followed by washing the enzyme with sample buffer (20 mM Tris-HCl, 10% glycerol and 10 mM sodium nitrate, pH 7.6) using an Amicon® Ultra Centrifugal Filter 30 MWCO. Next, both enzyme samples were diluted to 1 μM in aerobic sample buffer with 1M sodium bicarbonate (final pH of 7.9) to a final volume of 300 μL. These samples were placed in closed flasks with mineral oil occupying the volume of the headspace (150 μL). The same procedure was followed for the control, using anaerobic buffer with bicarbonate. To check for product inhibition during these experiments, an additional control was performed with WT *Dv*FdhAB where the activity was measured for CO_2_ reduction. For each timepoint, a sample was collected, and formate oxidation or CO_2_ reduction activities were measured as described.^24^

For the assays with superoxide dismutase (SOD) and catalase a similar procedure was followed, with small changes. WT *Dv*FdhAB was diluted in aerobic sample buffer with 20 mM sodium formate and without sodium nitrate. Four samples were prepared: (i) 1 μM *Dv*FdhAB, (ii) 1 μM *Dv*FdhAB and 100 U SOD, (iii) 1 μM *Dv*FdhAB and 4 nM catalase and (iv) 1 μM *Dv*FdhAB, 100 U SOD and 4 nM catalase. In samples (ii), (iii) and (iv), *Dv*FdhAB was the last enzyme added to the mixtures. The flasks were closed and incubated for 30 mins. The control assay was performed with WT *Dv*FdhAB diluted in anaerobic sample buffer.

The kinetic assays under aerobic conditions were performed using WT *Dv*FdhAB and the C872A variant, which is equivalent to the active form of *Dv*FdhAB.^25^ The experiments were performed similarly to what is described for FdhAB anaerobic assays,^24^ except for the redox mediators used. In these experiments, 1 mM PMS and 100 μM DCPIP were used, instead of 2 mM BV. The reduction of DCPIP by PMS was followed at 600 nm (ε_600_(DCPIP) = 20.7 mM^-1^ cm^-1^). As control anaerobic activity was also determined using PMS and DCPIP.

### Thermal Shift Assays

The Thermal Shift Assays were performed using the StepOnePlus System from Applied Biosystems with an excitation wavelength of 470 nm and the ROX fluorescence emission filter (≈ 610nm). *Dv*FdhAB (1 µM) was incubated with and without sodium formate (10 mM) for 10 minutes at room temperature and submitted to the assay in 250 mM potassium phosphate pH 7.0 with the Protein Thermal Shift Dye (1X, Applied Biosystems). Fluorescence was monitored and unfolding curves were generated using a temperature gradient from 25 to 95 °C in 46 min. All experiments were performed in triplicate, and the reported T_m_ values are based on the mean values determined from the minimum value of the inverse of the first derivative of the experimental data.

### NMR experiments

NMR data were acquired at room temperature (∼285K) on a 500 MHz Bruker NEO spectrometer (Bruker, Wissembourg, France) equipped with a 5 mm inverse detection triple-resonance z-gradient probe head (TXI) and processed using the software TopSpin 4.2.0 (Bruker BioSpin). All samples were prepared in 90%H_2_O + 10%D_2_O, 20 mM Tris-HCl buffer, pH 8 with 150 mM NaCl and 100 μM 4,4-dimethyl-4-silapentanesulfonic acid (DSS), used as a chemical shift standard. Three samples were prepared where the concentration of protein was kept at 0.5 µM and the concentration of ligand (formate) was 1, 10 and 50 mM. A fourth sample with 10 mM formate, was also prepared in anaerobic conditions. For each sample, a reference was also prepared in the same conditions but lacking the protein. All 1H spectra were acquired in a spectral window of 8196.72 Hz centered at 2354.10 Hz with 8 transients, 32 K data points, and a relaxation delay of 1.0 s. The solvent suppression was performed using an excitation sculpting scheme with gradients^47^ in which the solvent signal was irradiated with a selective pulse (Squa100.1000) with a length of 2 ms. The time for each experiment was 36s. For each sample, 65 1H spectra were acquired, corresponding to a total of ∼40 min of reaction time.

### EPR experiments

EPR analysis was performed on a Bruker ELEXSYS E500 spectrometer equipped with an ER4102ST standard rectangular Bruker EPR cavity fitted to an Oxford Instruments EPR 900 helium flow cryostat. A sample of 0.5 mL of 100 µM WT *Dv*FdhAB enzyme in MOPS buffer was degassed in the glove box and incubated with 10 mM formate for 10 min. A first EPR sample was taken and frozen anaerobically in liquid nitrogen. Then, the enzyme solution was left under air for more than 2 hours for oxidation and a second EPR sample was taken and frozen.

The enzyme solutions were then degassed again in the glove box and incubated with formate for reduction. A third EPR sample was taken and frozen anaerobically. After EPR analysis, this third sample was anaerobically thawed in the glove box and additionally reduced with 10 mM dithionite. A second set of similar experiments was performed to compare with much longer incubation time with formate (70 h) and oxygen treatment (20 h under air). All samples were studied by EPR at 15K and 80 K to analyze the behavior of the FeS centers and the W cofactor, respectively. Spin intensity measurements were performed by double integration of EPR spectra recorded in non-saturating conditions and comparison with a 1 mM Cu(II)EDTA standard.

### Computational calculations of magnetic parameters

The structural models of the tungsten cofactor were created using the atomic coordinates from the Reox_120min structure (PDB ID: 8RC9). The geometry of the models was optimized under two constraints: (i) the SSSS dihedral angle of the two dithiolates has been fixed to -33°, which correspond to the value in Reox_120min; (ii) the positions of the tungsten and the alpha carbon of the selenocysteine have been frozen. All calculations were performed at a DFT level of theory using the B3LYP functional with the D3BJ dispersion correction.^48^ The def2-SVP basis set was chosen with the RI approximation for the geometry optimization. The determination of EPR parameters was carried out using the segmented all-electron relativistically contracted (SARC) basis set of triple-ζ quality (TZVP) for tungsten and the def2-TZVPP basis set for the other atoms.^49^ The relativistic effect was described by the ZORA method^50^ and spin-orbit couplings were considered with the SOMF treatment. All DFT calculations were performed using Orca 5.0.3 program package.^51^

The *Dv*FdhAB structures obtained in this work are deposited in the Protein Data Bank, under accession codes: 8RC8, 8RC9, 8RCA, 8RCB and 8RCC. Tables and Figures explaining more extensively the results described in this article and additional Figures presenting structural superpositions (Supplementary Material (PDF)).

### Funding Sources

This work was financially supported by Fundação para a Ciência e Tecnologia (FCT, Portugal) through fellowship nos. 2023.00286.BD (G.V.-A.) and DFA/BD/7897/2020 (R.R.M.), grant no. PTDC/BII-BBF/2050/2020 (http://doi.org/10.54499/PTDC/BII-BBF/2050/2020) (I.A.C.P. and M.J.R.), research contract 2020.00043.CEECIND (A.V.) and R&D units MOSTMICRO-ITQB (grant nos. UIDB/04612/2020 and UIDP/04612/2020) (I.A.C.P.) and UCIBIO (grant nos. UIDP/04378/2020 and UIDB/04378/2020) (M.J.R.), and Associated Laboratories LS4FUTURE (grant no. LA/P/0087/2020) (I.A.C.P.) and i4HB (grant no. LA/P/0140/2020) (M.J.R.). The NMR spectrometers at FCT-NOVA are part of Rede Nacional de RMN (PTNMR), supported by FCT-MCTES (ROTEIRO/0031/2013-PINFRA/22161/2016) (cofinanced by FEDER through COMPETE 2020, POCI and PORL and FCT through PIDDAC). This work was also funded by the French national research agency (ANR – MOLYERE project, grant no. 16-CE-29-0010-01) (B.G.) and supported by the computing facilities of the Centre Régional de Compétences en Modélisation Moléculaire de Marseille).

## Acknowledgments

We thank Prof Carlos Romão for insightful discussions pertaining the coordination chemistry of the SeCys unbound active site and the excellent technical assistance of João Carita from ITQB NOVA on microbial cell growth. We are also grateful to the EPR-MRS facilities of the Aix-Marseille University EPR centre and acknowledge the support of the European research infrastructure MOSBRI (grant no. 101004806) (B.G.) and the French research infrastructure INFRANALYTICS (FR2054) (B.G.). We also acknowledge the ESRF Synchrotron for provision of synchrotron radiation facilities, and we thank the staff of the ESRF and EMBL Grenoble for assistance and support in using the HPMX lab and beamlines ID23-1, ID30A-3, ID30B.

## Abbreviations

AOR: Aldehyde Oxido-reductase
DTT: Dithiothreitol
*Dv*: *Desulfovibrio vulgaris*
EPR: Electron Paramagnetic Resonance
Fdh: Formate dehydrogenase
HP: High Pressure
MGD: Molybdopterin Guanine Dinucleotides
ND: New drop
ROS: Reactive Oxygen Species
SOD: Superoxide dismutase
TSA: Thermal Shift Assay

