## Supplementary Material for "Substrate-dependent oxidative inactivation of a W-dependent formate dehydrogenase involving selenocysteine displacement"

### Index

**Table S 1- Description of the procedure used to obtain the different structures of DvFdhAB in this and other works.**

| Structure | Procedure |
|---|---|
| As-isolated [1]<br>PDB_ID: 6SDR | *Dv*FdhAB crystallized **as-isolated** in the presence of $O_2$ (As-isolated). |
| Formate-reduced [1]<br>PDB_ID: 6SDV | *Dv*FdhAB co-crystallized with 10 mM of sodium formate and **not exposed** to $O_2$ (Reduced). |
| Control_Red<br>PDB_ID: 8RC8 | *Dv*FdhAB co-crystallized with 10 mM of sodium formate and **not exposed** to $O_2$ (Reduced). |
| Reox_12min [2]<br>PDB_ID: 8BQL | *Dv*FdhAB co-crystallized with 10 mM of sodium formate; plate well opened and crystals exposed to atmospheric $O_2$, for **12 min**, while still in the original drop (which **contains** 10 mM of sodium formate), and then flash cooled in liquid nitrogen (as-isolated/Oxidized). |
| Reox_120min<br>PDB_ID: 8RC9 | *Dv*FdhAB co-crystallized with 10 mM of sodium formate; plate well opened and the crystals were exposed to atmospheric $O_2$, for **120 min**, while still in the original drop (which **contains** 10 mM of sodium formate), and then flash cooled in liquid nitrogen (W-O=O…SeCys form). |
| Reox_ND_NoFormate<br>PDB_ID: 8RCA | *Dv*FdhAB co-crystallized with 10 mM of sodium formate; the crystals were transferred from the original drop to a **N**ew **D**rop (oxygenated), mimicking the mother liquor but **without** sodium formate, and were exposed to atmospheric $O_2$, for 60 min, and then flash cooled in liquid nitrogen (as-isolated/Oxidized). |
| Reox_ND_Formate<br>PDB_ID: 8RCB | *Dv*FdhAB co-crystallized with 10 mM of sodium formate; the crystals were transferred from the original drop to a **N**ew **D**rop (oxygenated), mimicking the mother liquor **containing** 10 mM of sodium formate, and were exposed to atmospheric $O_2$, for 34 min, and then flash cooled in liquid nitrogen (W-O=O…SeCys form). |
| HP_$CO_2$<br>PDB_ID: 8RCC | *Dv*FdhAB aerobically crystallized **without** sodium formate; the crystals were pressurised with 48 bar of $CO_2$, and then flash cooled in liquid nitrogen, under 200 bar helium (W-O=O…SeCys form). |

**Table S 2- Distances between the W ion, Se atom of SeCys and the two oxygen atoms of the dioxygen molecule for the Reox_120min, Reox_ND_Formate and HP_$CO_2$ structures and respective resolutions.**

| Distance (Å) | Reox_120min (2.06 Å) | Reox_ND_Formate (2.11 Å) | HP_$CO_2$ (2.30 Å) |
|---|---|---|---|
| W - - - O1 | 2.34 | 2.32 | 2.43 |
| W - - - O2 | 2.52 | 2.64 | 1.93 |
| W - - - Se | 4.20 | 4.48 | 3.69 |
| O1 - - - Se | 2.57 | 3.01 | 2.89 |
| O2 - - - Se | 2.46 | 2.62 | 2.19 |

**Table S 3- B-factor values for the W ion, sulfido ligand (S), the four sulfur atoms of the 2 dithiolenes, Se atom of SeCys and the two atoms of the dioxygen molecule for the Control_Red, Reox_12min, Reox_120min, Reox_ND_No_Formate, Reox_ND_Formate and HP_$CO_2$ structures and respective resolutions.**

| **B-factor ($Å^2$)** | **Control_Red (2.00 Å)** | **Reox_12min (1.91 Å)** | **Reox_120min (2.06 Å)** | **Reox_ND_No_Formate (1.66 Å)** | **Reox_ND_ Formate (2.11 Å)** | **HP_$CO_2$ (2.30 Å)** |
|---|---|---|---|---|---|---|
| **W** | 24.09 | 24.75 | 35.67 | 21.05 | 33.75 | 30.77 |
| **S** | 28.66 | 26.61 | 58.88 | 18.74 | 60.37 | 43.89 |
| **S12**$_{MGD1}$ | 22.13 | 21.90 | 29.10 | 21.74 | 24.24 | 26.59 |
| **S13**$_{MGD1}$ | 21.80 | 24.55 | 30.23 | 21.88 | 24.30 | 27.26 |
| **S12**$_{MGD2}$ | 24.31 | 26.19 | 32.19 | 21.22 | 37.30 | 29.81 |
| **S13**$_{MGD2}$ | 23.65 | 25.26 | 36.35 | 20.72 | 29.61 | 26.54 |
| **Se** | 25.54 | 29.30 | 53.55 | 25.10 | 92.48 | 75.80 |
| **Cβ**$_{U192}$ | 27.58 | 30.10 | 49.66 | 24.37 | 55.09 | 56.78 |
| **Cα**$_{U192}$ | 28.29 | 29.85 | 44.20 | 24.09 | 40.43 | 49.87 |
| **O1** | NA | NA | 44.42 | NA | 25.44 | 32.52 |
| **O2** | NA | NA | 32.91 | NA | 39.70 | 33.80 |

**Table S 4- Crystallographic data processing and refinement statistics.**

| **Crystal** | **Control_Red** | **Reox_120min** | **Reox_120min**_Staraniso |
|---|---|---|---|
| **$PDB_{code}$** | 8RC8 | | 8RC9 |
| **Diffraction Data** | | | |
| Space group | $P2_12_12_1$ | $P2_12_12_1$ | $P2_12_12_1$ |
| Cell dimensions (Å) (º) | a=64.99, b=124.35, c=149.90 | a=65.18, b=127.93, c=149.47 | a=65.18, b=127.93, c=149.47 |
| Wavelength (Å) | 0.9686 | 0.9686 | 0.9686 |
| Beamline | ESRF ID30B | ESRF ID30B | ESRF ID30B |
| No. Crystals | 1 | 1 | 1 |
| Resolution range of data (Å) (last shell) | 47.85 – 2.00 (2.04 – 2.00) | 97.19 – 2.28 (2.32 – 2.28) | 97.19 – 2.06 (2.15 – 2.06) |
| Completeness (%) (last shell) | 99.60 (92.40) | 96.52 (98.31) | 87.87 (49.15) |
| Rmerge (last shell) | 0.182 (1.224) | 0.110 (0.658) | 0.119 (1.110) |
| Rmeas (last shell) | 0.214 (1.429) | 0.127 (0.758) | 0.137 (1.241) |
| I/σI (last shell) | 6.6 (1.3) | 7.9 (2.1) | 7.0 (1.5) |
| CC 1/2 (last shell) | 0.994 (0.527) | 0.991 (0.632) | 0.991 (0.495) |
| Redundancy (last shell) | 7.0 (6.6) | 4.1 (4.2) | 4.3 (5.1) |
| **Refinement** | | | |
| Reflections used in refinement (work (free)) | 77848 (4152) | | 65087 (3244) |
| Rwork | 0.183 | | 0.206 |
| Rfree | 0.219 | | 0.255 |
| Nº of non-hydrogen atoms | 9804 | | 9587 |
| Protein | 9205 | | 9224 |
| Ligands | 175 | | 154 |
| Ions | 2 | | 2 |
| Solvent | 422 | | 207 |
| **Geometry and *B*-factors** | | | |
| RMSD bond lengths (Å) | 0.004 | | 0.005 |
| RMSD bond angles (º) | 1.052 | | 1.393 |
| Average B-factor ALL ($Å^2$) | 31.24 | | 40.72 |
| Protein ($Å^2$) | 33.28 | | 43.64 |
| Ligands ($Å^2$) | 31.06 | | 34.28 |
| Ions ($Å^2$) | 26.38 | | 47.27 |
| Solvent ($Å^2$) | 33.34 | | 33.82 |
| Ramachandran favoured (%) | 96.32 | | 95.47 |
| Ramachandran outliers (%) | 0.17 | | 0.34 |
| Molprobity score | 1.26 | | 1.50 |
| Clashscore | 2.38 | | 2.98 |

| **Crystal** | **Reox_ND_NoFormate** | **Reox_ND_Formate** | **Reox_ND_Formate**_Staraniso |
|---|---|---|---|
| **PDB$_{code}$** | 8RCA | | 8RCB |
| **Diffraction Data** | | | |
| Space group | $P2_12_12_1$ | $P2_12_12_1$ | $P2_12_12_1$ |
| Cell dimensions (Å) (º) | a=64.56, b=128.01, c=148.91 | a=64.43, b=123.09, c=148.14 | a=64.43, b=123.09, c=148.14 |
| Wavelength (Å) | 0.7749 | 0.9686 | 0.9686 |
| Beamline | ESRF ID23-1 | ESRF ID30B | ESRF ID30B |
| No. Crystals | 1 | 1 | 1 |
| Resolution range of data (Å) (last shell) | 97.26 – 1.66 (1.79 – 1.66) | 94.67 – 2.42 (2.46 – 2.42) | 94.67 – 2.11 (2.28 – 2.11) |
| Completeness (%) (last shell) | 90.60 (53.30) | 98.64 (98.80) | 91.15 (40.48) |
| Rmerge (last shell) | 0.104 (1.098) | 0.140 (1.070) | 0.153 (1.187) |
| Rmeas (last shell) | 0.117 (1.201) | 0.149 (1.132) | 0.163 (1.279) |
| I/σI (last shell) | 7.9 (1.5) | 10.4 (2.1) | 8.4 (1.6) |
| CC 1/2 (last shell) | 0.996 (0.658) | 0.996 (0.774) | 0.996 (0.591) |
| Redundancy (last shell) | 4.66 (6.12) | 9.0 (9.4) | 8.9 (7.3) |
| **Refinement** | | | |
| Reflections used in refinement (work (free)) | 102948 (5448) | | 54095 (2723) |
| Rwork | 0.196 | | 0.199 |
| Rfree | 0.232 | | 0.240 |
| Nº of non-hydrogen atoms | 9971 | | 9410 |
| Protein | 9228 | | 9153 |
| Ligands | 172 | | 161 |
| Ions | 2 | | 2 |
| Solvent | 569 | | 94 |
| **Geometry and *B*-factors** | | | |
| RMSD bond lengths (Å) | 0.008 | | 0.003 |
| RMSD bond angles (º) | 1.511 | | 1.212 |
| Average B-factor ALL (Å$^2$) | 30.65 | | 38.68 |
| Protein (Å$^2$) | 32.60 | | 41.18 |
| Ligands (Å$^2$) | 25.27 | | 33.93 |
| Ions (Å$^2$) | 19.89 | | 47.06 |
| Solvent (Å$^2$) | 33.35 | | 28.54 |
| Ramachandran favoured (%) | 96.33 | | 95.85 |
| Ramachandran outliers (%) | 0.26 | | 0.17 |
| Molprobity score | 1.54 | | 1.20 |
| Clashscore | 4.43 | | 1.63 |

| **Crystal** | **HP_$CO_2$** | **HP_$CO_2$**_Staraniso |
|---|---|---|
| **PDB$_{code}$** | | 8RCC |
| **Diffraction Data** | | |
| Space group | $P2_12_12_1$ | $P2_12_12_1$ |
| Cell dimensions (Å) (º) | a=64.81, b=124.40, c=150.31 | a=64.81, b=124.40, c=150.31 |
| Wavelength (Å) | 0.9677 | 0.9677 |
| Beamline | ESRF ID30A-3 | ESRF ID30A-3 |
| No. Crystals | 1 | 1 |
| Resolution range of data (Å) (last shell) | 95.84 – 2.66 (2.70 – 2.66) | 95.84 – 2.30 (2.44 – 2.30) |
| Completeness (%) (last shell) | 96.26 (99.05) | 89.19 (45.52) |
| Rmerge (last shell) | 0.194 (0.901) | 0.239 (1.421) |
| Rmeas (last shell) | 0.214 (0.995) | 0.264 (1.546) |
| I/σI (last shell) | 8.1 (2.1) | 6.5 (1.4) |
| CC 1/2 (last shell) | 0.986 (0.639) | 0.985 (0.465) |
| Redundancy (last shell) | 5.5 (5.5) | 5.6 (6.3) |
| **Refinement** | | |
| Reflections used in refinement (work (free)) | | 42746 (2267) |
| Rwork | | 0.194 |
| Rfree | | 0.244 |
| Nº of non-hydrogen atoms | | 9544 |
| Protein | | 9193 |
| Ligands | | 246 |
| Ions | | 2 |
| Solvent | | 103 |
| **Geometry and *B*-factors** | | |
| RMSD bond lengths (Å) | | 0.005 |
| RMSD bond angles (º) | | 1.360 |
| Average B-factor ALL (Å$^2$) | | 33.54 |
| Protein (Å$^2$) | | 35.93 |
| Ligands (Å$^2$) | | 35.28 |
| Ions (Å$^2$) | | 37.33 |
| Solvent (Å$^2$) | | 25.44 |
| Ramachandran favoured (%) | | 94.86 |
| Ramachandran outliers (%) | | 0.34 |
| Molprobity score | | 1.93 |
| Clashscore | | 5.17 |

**Table S 5- DFT-predicted g-values for the optimized $W^V$ structural models (Figure S11).**

| Model | Calculated g-values |
|---|---|
| 1 | 2.189 2.049 2.026 |
| 2 | 2.019 2.013 2.013 |
| 3 | 2.616 2.171 2.057 |
| 4 | n.d. [1] |
| 5 | n.d. [1] |
| 6 | n.d. [1] |

[1] for these models, the optimized structure cannot fit with the rest of the protein due to strong distortions of dithiolene moieties.

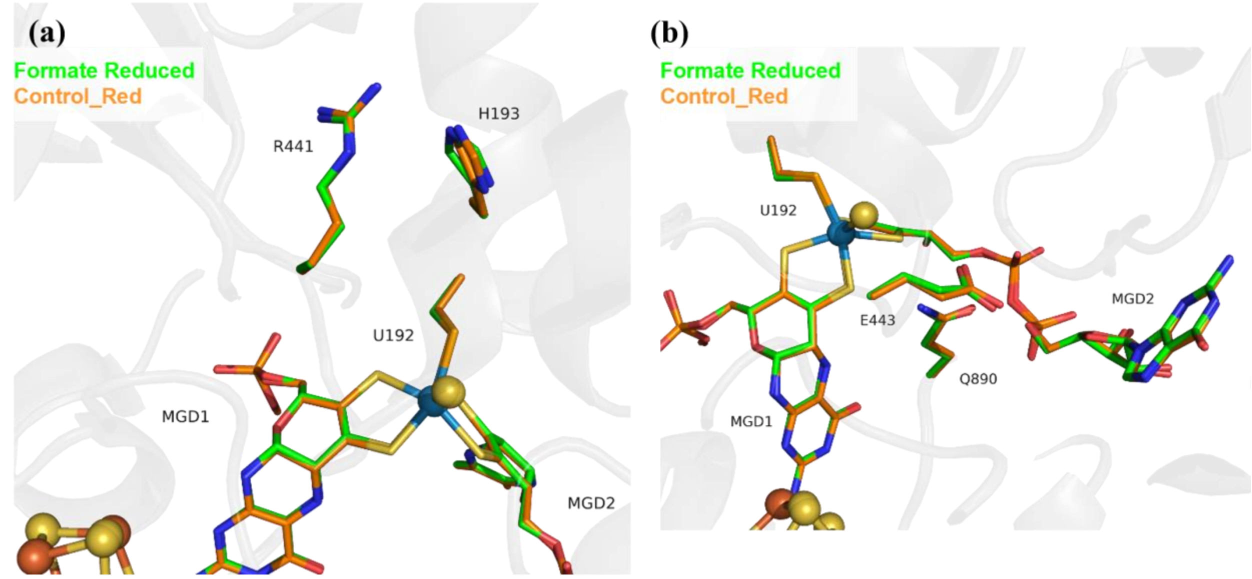

**Figure S 1- Superposition of *Dv*FdhAB WT formate reduced (PDB_ ID: 6SDV) (green) and Control_Red structure (orange) in two different views.**
Rmsd is 0.21 Å for 964 Cα atoms of *Dv*FdhA, and of 0.19 Å for 214 Cα atoms of *Dv*FdhB. **(a)** U192, H193, R441 and the two MGD co-factors are shown as sticks. **(b)** U192, E443, Q890 and the two MGD co-factors are shown as sticks.

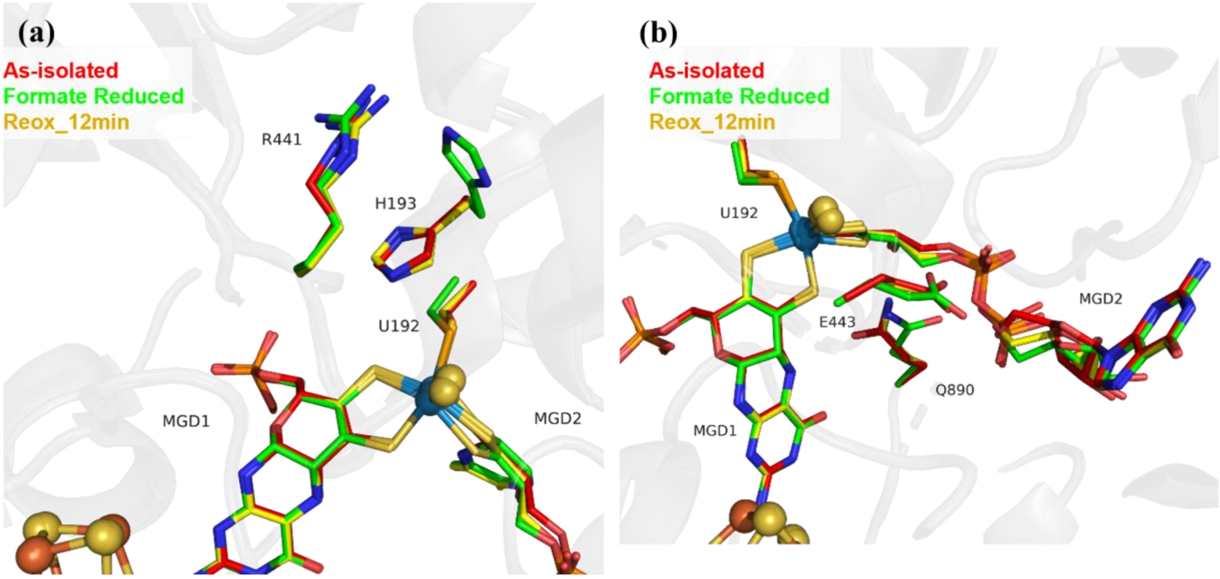

**Figure S 2- Superposition of *Dv*FdhAB WT WT as-isolated (oxidized) (PDB_ID: 6SDR) (red), formate reduced (green) and oxygen exposed Reox_12min structure (yellow) in two different views.** Rmsd is 0.15 Å for 963 Cα atoms of *Dv*FdhA, and of 0.17 Å for 214 Cα atoms of *Dv*FdhB. **(a)** U192, H193, R441 and the two MGD co-factors are shown as sticks. **(b)** U192, E443, Q890 and the two MGD co-factors are shown as sticks.

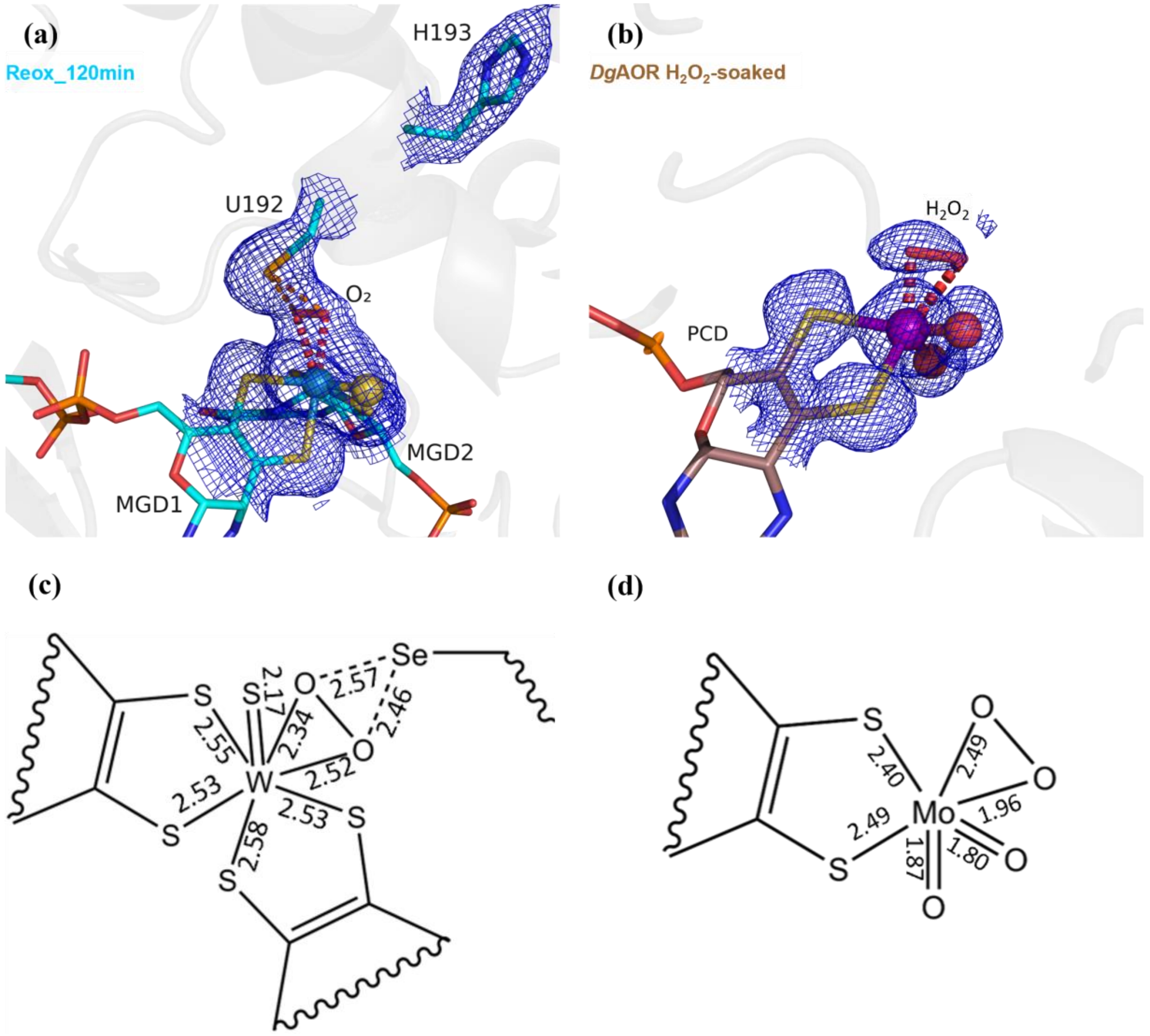

**Figure S 3- Comparison of the metal coordination between the dioxygen/peroxide containing active sites of DvFdhAB Reox_120min (cyan) and DgAOR H2O2-soaked (PDB_ID: 4C80) (brown).**

(a) DvFdhAB Reox_120min structure (cyan) and respective electron density map (2Fo-Fc), at 1σ (blue mesh). U192, H193, the two MGD co-factors coordinating the W ion and the dioxygen molecule (red) are shown as sticks. (b) DgAOR H2O2-soaked (PDB_ID: 4C80) structure (brown) and respective electron density map (2Fo-Fc), at 1σ (blue mesh). The Molybdopterin Cytosine Dinucleotide (MCD) co-factor coordinating the Mo ion, the two terminal oxo ligands and peroxide molecule (red) are shown as sticks. (c) 2D representation of the DvFdhAB Reox_120min W active site, indicating the bond lengths (in Å) of relevant bonds. (d) 2D representation of the DgAOR H2O2-soaked (PDB_ID: 4C80) Mo active site, indicating the bond lengths (in Å) of relevant bonds.

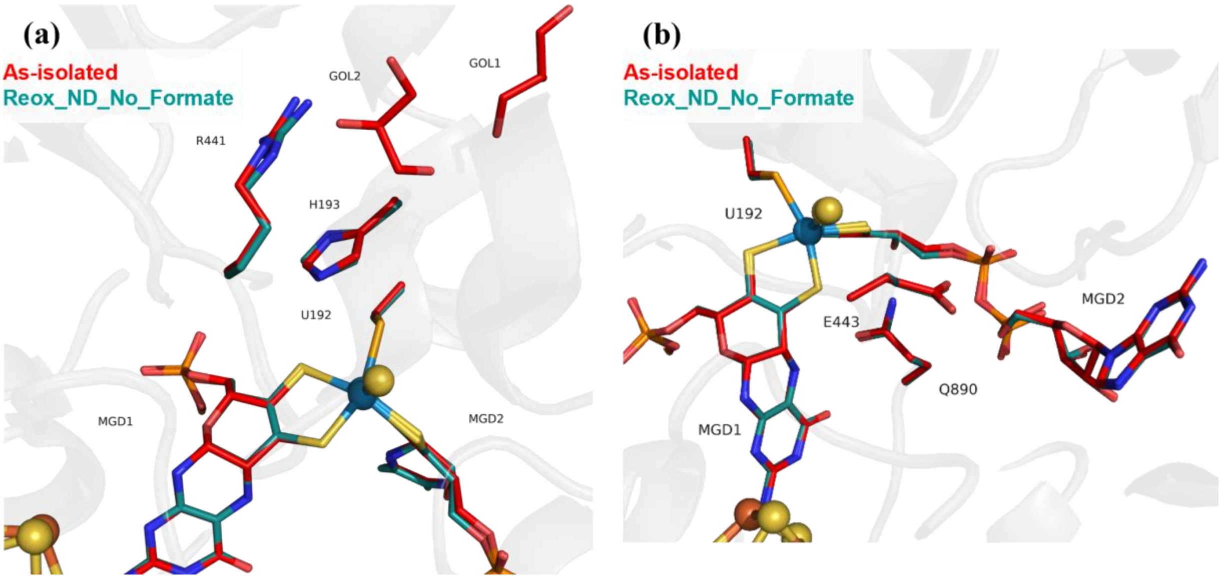

**Figure S 4- Superposition of DvFdhAB WT as-isolated (oxidized) (PDB_ID: 6SDR) (red) and oxygen exposed Reox_ND_NoFormate structure (blue) in two different views.**
Rmsd is 0.15 Å for 963 Cα atoms of DvFdhA, and of 0.15 Å for 214 Cα atoms of DvFdhB. (a) U192, H193, R441, the two MGD co-factors and two glycerol molecules are shown as sticks. (b) U192, E443, Q890 and the two MGD co-factors are shown as sticks.

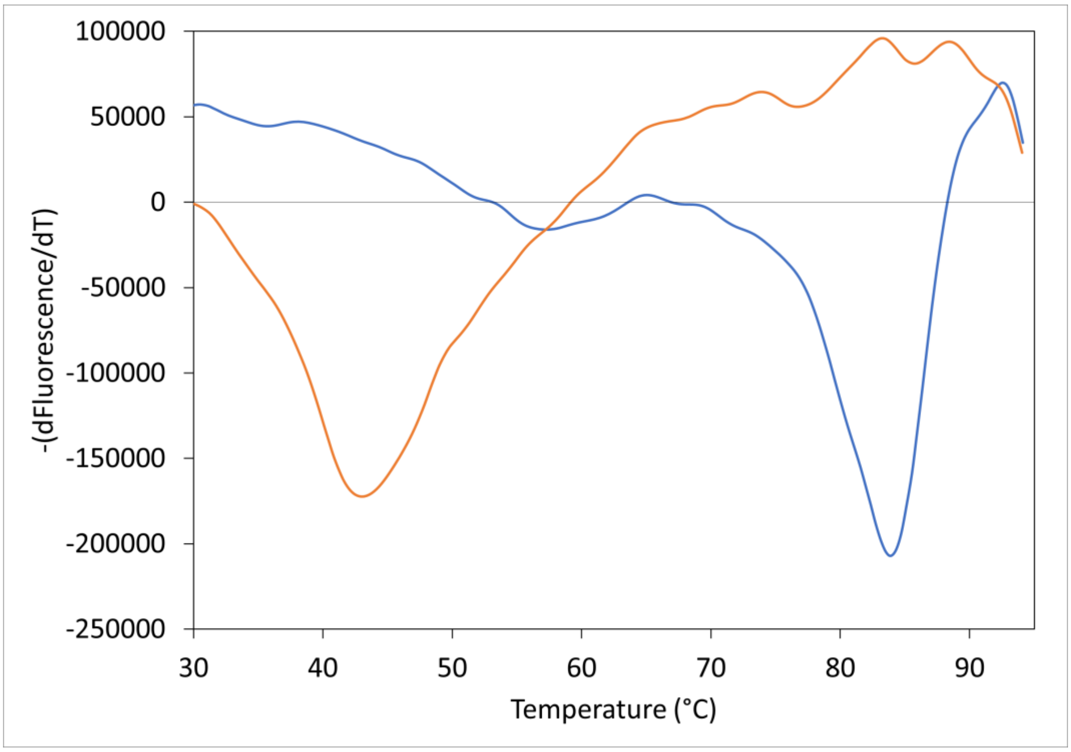

**Figure S 5- *Dv*FdhAB WT Thermal Shift Assay in the presence and absence of formate.**
The inverse of the derivative of fluorescence with respect to time is plotted as a function of the temperature for the assays with 10 mM of formate (orange line) and without (blue line). The calculated $T_m$ are 83,8 ± 0.2 ˚C and 43.2 ± 0.2 ˚C for *Dv*FdhAB WT in the absence and presence of 10 mM of formate, respectively.

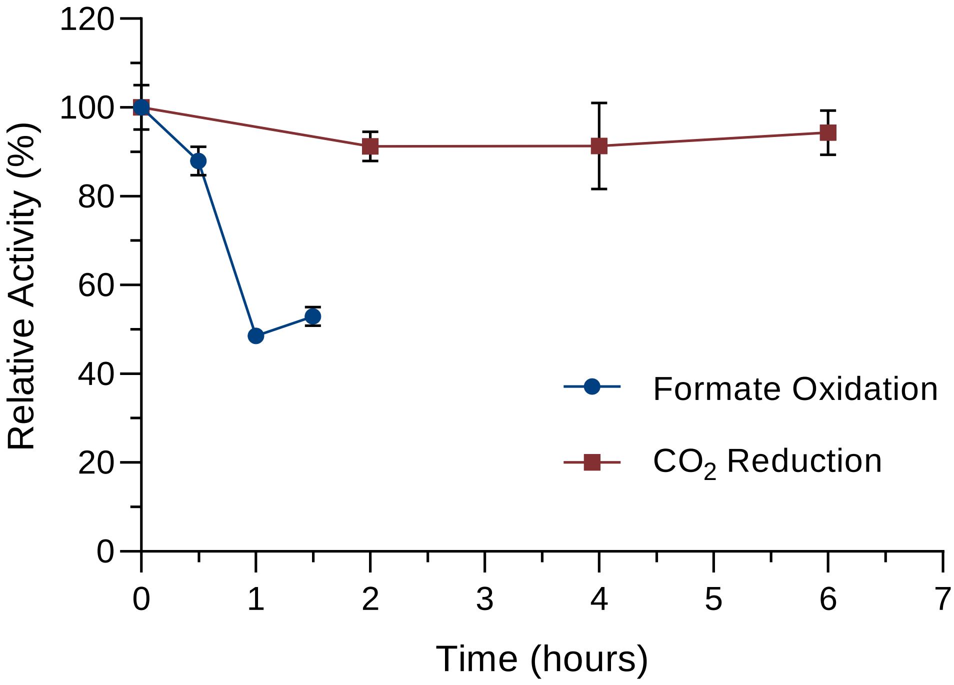

**Figure S 6- Relative activity for $CO_2$ reduction (red) and formate oxidation (blue) of as-isolated *Dv*FdhAB during oxygen exposure in the presence of $CO_2$.**
T=0h was considered as 100% of activity. Data are presented as mean values ± s.d. (n = 3 assay technical replicates).

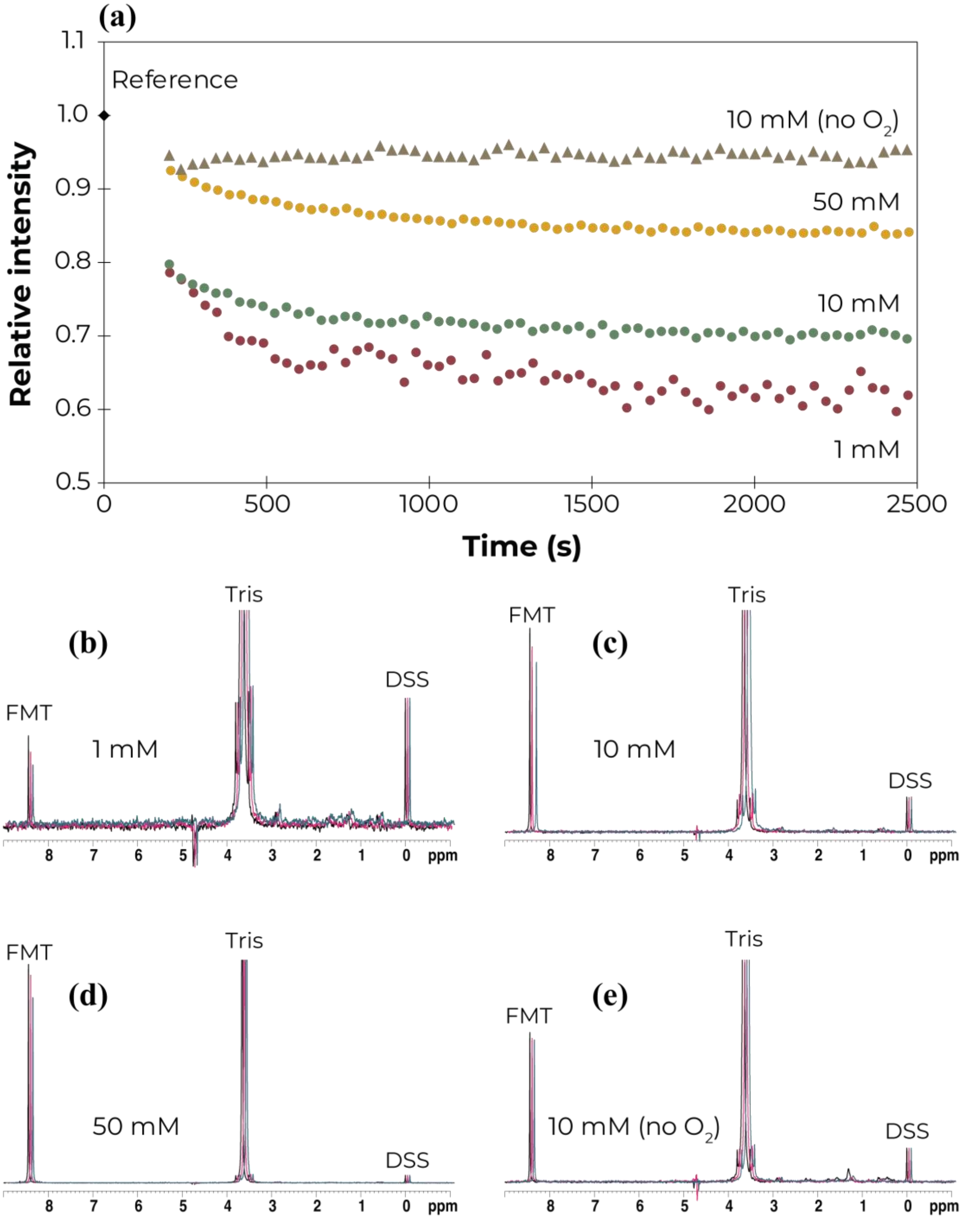

**Figure S 7- DvFdhAB activity assays by 1H NMR.**
(a) The relative intensity of the 1H NMR formate peak (calculated as the intensity ratio between the formate peak and that of DSS) is plotted as a function of time for the activity assays with formate concentrations of 1 mM (red circle), 10 mM (green circle), 50 mM (yellow circle); and 10 mM in the absence of atmospheric oxygen (grey triangle), the reference (which is calculated in the same way as before, but using the samples prepared in the absence of enzyme) is shown (black diamond). (b - e) Representative 1D 1H spectra of each condition tested (1, 10 and 50 mM formate and 10 mM formate in the absence of atmospheric oxygen, respectively) (black: reference 1H spectrum; magenta: 1st 1H spectrum; blue: last 1H spectrum). Above each spectra the peaks of FMT (formate), Tris (buffer) and DSS are indicated. The spectra corresponding to the 1st and last acquisition points are shifted to the right for better analysis.

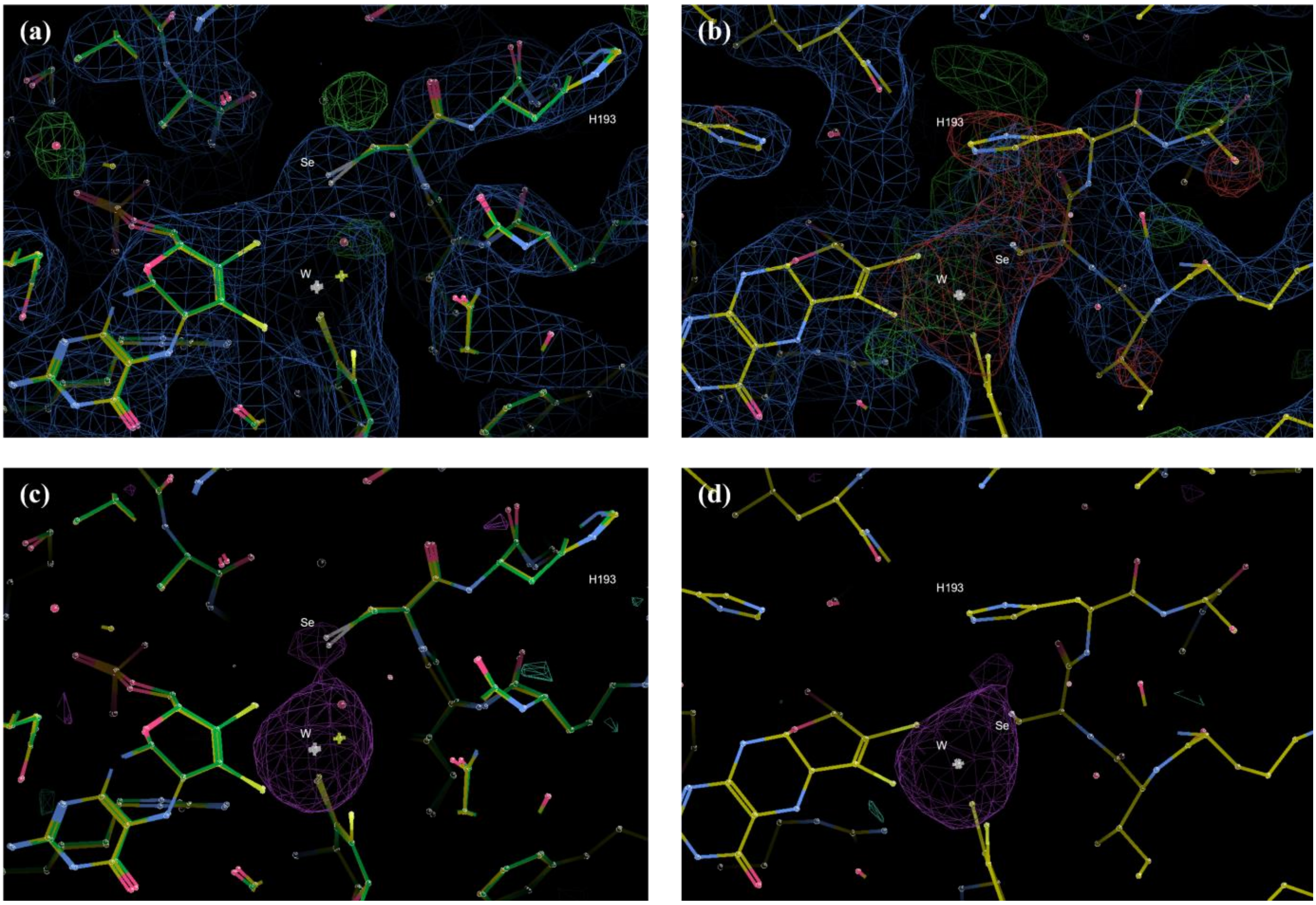

**Figure S 8- Solved, unrefined, structure (yellow) obtained from the *Dv*FdhAB WT crystals produced with the concentrated solution left from the NMR experiments with formate under air, at a resolution of 2.83 Å.** 2Fo-Fc maps at 1 σ (blue mesh), Fo-Fc maps at 3 σ (green and red mesh, respectively for positive and negative densities) and anomalous map peaks at 3 σ (violet mesh) are shown. Images produced with Coot [3]. (a) and (c) *Dv*FdhAB WT "after-NMR" structure, solved using Reox_120min as molecular replacement model (yellow), superposed with the Reox_120min structure (green). (b) and (d) *Dv*FdhAB WT "after-NMR" structure, solved using *Dv*FdhAB WT as-isolated (oxidized) (PDB_ID: 6SDR) as molecular replacement model (yellow).

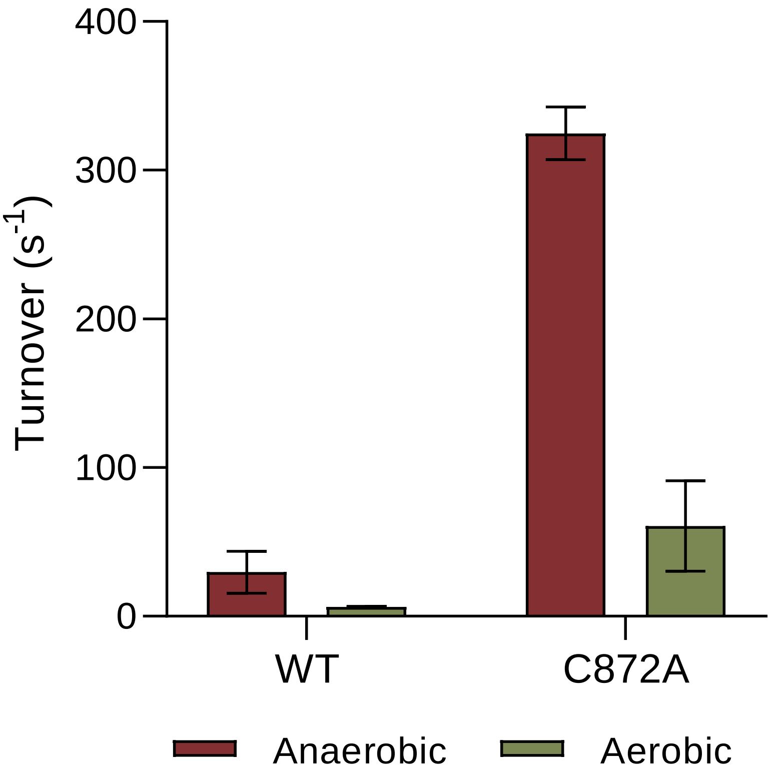

**Figure S 9- Formate oxidation activity of as-isolated WT *Dv*FdhAB and C872A variant (this variant that is equivalent to the DTT-activated form [4]).**
With PMS + DCPIP as artificial electron acceptors in anaerobic conditions (dark red, glove box) and in aerobic conditions (green). Data are presented as mean values ± s.d. (n = at least 3 assay technical replicates). No DTT was used in the assays. The higher error observed for the aerobic assay of the C872A variant is due to the decreasing activity of the enzyme in these conditions.

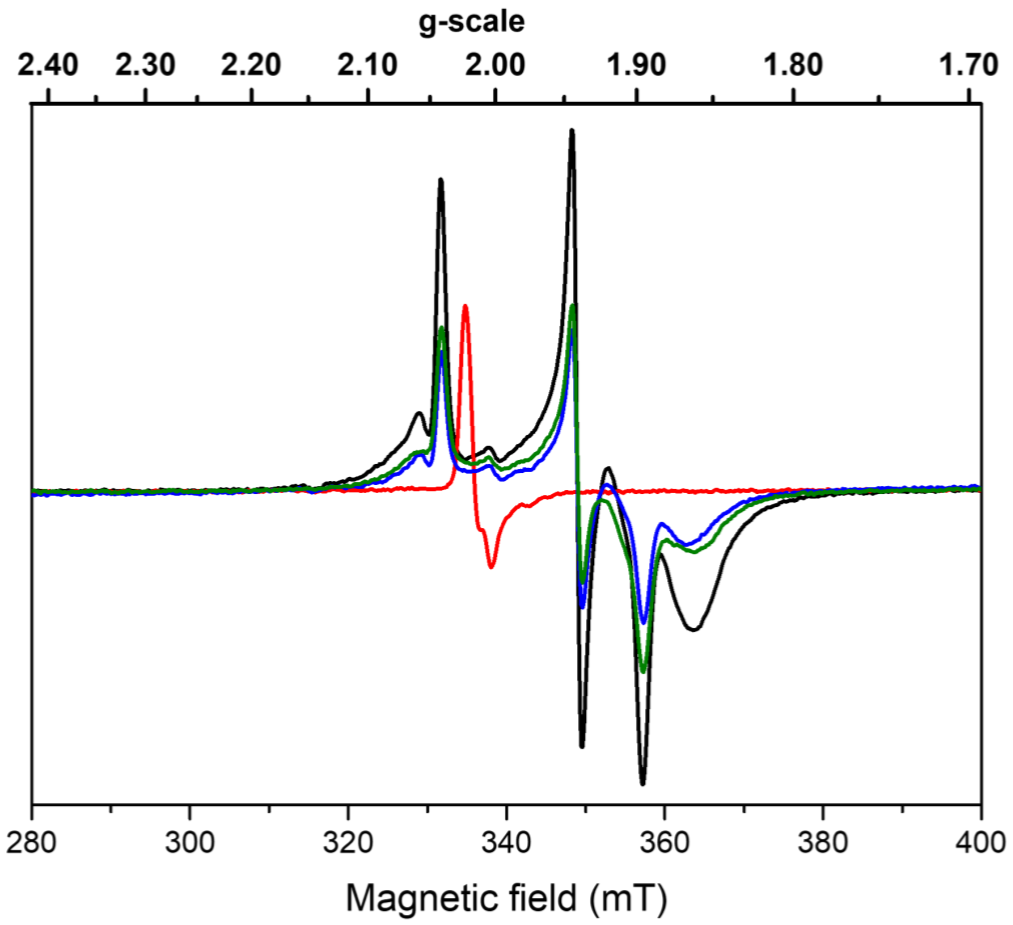

**Figure S 10- Influence of formate and oxygen exposure on FeS center EPR signals of *Dv*FdhAB.**
Anaerobic reduction with formate (black trace) followed by oxygen treatment (red trace), then degassing and anerobic reduction with formate (blue trace), and subsequent reduction with dithionite (green trace). EPR conditions: Temperature, 15 K; microwave power 1 mW at 9.479 GHz, modulation amplitude 1 mT at 100 kHz.

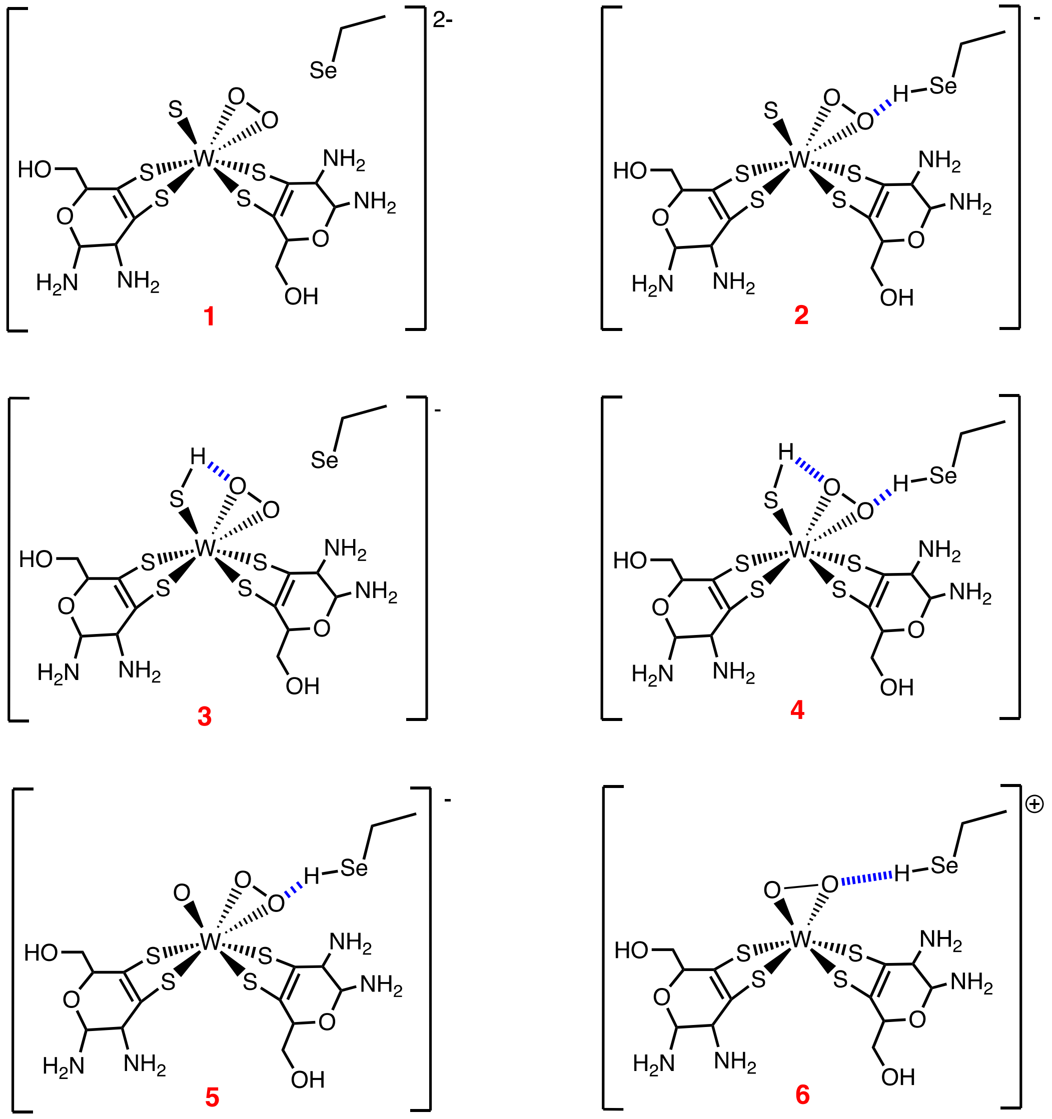

**Figure S 11- Structural models of the W cofactor used for DFT calculations.**
